# Targeting Acute Myeloid Leukemia Stem Cells Through Perturbation of Mitochondrial Calcium

**DOI:** 10.1101/2023.10.02.560330

**Authors:** Anagha Inguva Sheth, Krysta Engel, Hunter Tolison, Mark J Althoff, Maria L. Amaya, Anna Krug, Tracy Young, Shanshan Pei, Sweta B. Patel, Mohammad Minhajuddin, Amanda Winters, Regan Miller, Ian Shelton, Jonathan St-Germain, Tianyi Ling, Courtney Jones, Brian Raught, Austin Gillen, Monica Ransom, Sarah Staggs, Clayton A. Smith, Daniel A. Pollyea, Brett M. Stevens, Craig T. Jordan

**Affiliations:** Division of Hematology, University of Colorado School of Medicine, Aurora, CO, USA; Liangzhu Laboratory, Zhejiang University Medical Center, Bone Marrow Transplantation Center, Hangzhou, China; Division of Pediatric Hematology and Oncology, University of Colorado School of Medicine, Aurora, CO, USA; Princess Margaret Cancer Centre, University Health Network, Toronto, Canada; Department of Medical Biophysics, University of Toronto, Toronto, Canada

## Abstract

We previously reported that acute myeloid leukemia stem cells (LSCs) are uniquely reliant on oxidative phosphorylation (OXPHOS) for survival. Moreover, maintenance of OXPHOS is dependent on BCL2, creating a therapeutic opportunity to target LSCs using the BCL2 inhibitor drug venetoclax. While venetoclax-based regimens have indeed shown promising clinical activity, the emergence of drug resistance is prevalent. Thus, in the present study, we investigated how mitochondrial properties may influence mechanisms that dictate venetoclax responsiveness. Our data show that utilization of mitochondrial calcium is fundamentally different between drug responsive and non-responsive LSCs. By comparison, venetoclax-resistant LSCs demonstrate a more active metabolic (i.e., OXPHOS) status with relatively high steady-state levels of calcium. Consequently, we tested genetic and pharmacological approaches to target the mitochondrial calcium uniporter, MCU. We demonstrate that inhibition of calcium uptake sharply reduces OXPHOS and leads to eradication of venetoclax-resistant LSCs. These findings demonstrate a central role for calcium signaling in the biology of LSCs and provide a therapeutic avenue for clinical management of venetoclax resistance.

**Significance:** We identify increased utilization of mitochondrial calcium as distinct metabolic requirement of venetoclax-resistant LSCs and demonstrate the potential of targeting mitochondrial calcium uptake as a therapeutic strategy.

## Introduction

Targeting BCL-2 with the small molecule inhibitor, venetoclax, has become a mainstay in AML therapy. Multiple clinical studies have demonstrated that venetoclax (ven) in combination with hypomethylating agents (HMAs) including azacitidine and decitabine yields rates of complete remission (CR) that are comparable to conventional chemotherapy with considerably less toxicity, making the regimen particularly useful for elderly/unfit patients [1–4]. Despite the success of this regimen, approximately 30% of patients do not respond upfront and the majority of patients who initially achieve a response ultimately relapse on therapy [1, 3, 4]. Notably, while venetoclax-based regimens are able to directly target some LSCs [2], resistant LSCs either co-reside or evolve from sensitive LSCs [5, 6] and confer resistance. Consequently, there is an urgent need to develop therapeutic strategies to target venetoclax-resistant LSCs.

Regardless of sensitivity (or lack thereof) to venetoclax, a conserved property of all LSCs appears to be reliance on OXPHOS [5, 7–10]. While BCL-2 inhibition is able to target OXPHOS activity in sensitive LSCs, resistant LSCs display no change in metabolism upon venetoclax treatment [5, 9]. Consistent with this observation, multiple previous studies have demonstrated that various aspects of mitochondrial metabolism can be leveraged to overcome venetoclax resistance[11–14]. For example, one feature found in venetoclax-resistant AML cells in increased utilization of fatty acid oxidation (FAO) to drive OXPHOS. We have demonstrated that adding inhibition of FAO to venetoclax-based regimens is sufficient to restore inhibition of OXPHOS and consequently eradicate venetoclax-resistant LSCs [9]. Similarly, inhibition of MCL1 in the context of monocytic AML appears to function via an analogous mechanism and can also eradicate venetoclax-resistant LSCs [5, 9]. Another aspect of mitochondrial metabolism, increased reliance on NAD to drive OXPHOS, is also a therapeutic vulnerability, and represents yet another potential entry point for therapy[7] . Collectively, these studies overwhelmingly demonstrate the potential of manipulating LSC mitochondrial biology as an avenue to inhibit OXPHOS and thereby address venetoclax resistance.

While inhibiting OXPHOS in LSCs is an empirical goal, the initial molecular events that dictate venetoclax response remain elusive. Thus, in the present study, we investigated early-acting BCL-2 mediated pathways that may influence OXPHOS activity to determine if they are differentially utilized in sensitive versus resistant LSCs. One such pathway is BCL-2 mediated regulation of intracellular calcium dynamics [15]. It has been well established that BCL-2 modulates intracellular calcium localization by interacting with and activating or inhibiting calcium channels at the ER and mitochondria [15–19]. Further, mitochondrial calcium levels can directly influence OXPHOS activity as calcium is a cofactor for multiple metabolic enzymes [20, 21]. Calcium overload in the mitochondrial matrix has been shown to suppress OXPHOS activity by inhibiting the activity of key metabolic enzymes and electron transport chain activity [22–30]. Similarly, insufficient calcium can also reduce OXPHOS activity [31, 32]. Consequently, the role of mitochondrial calcium has been deemed as a “goldilocks” effect where either insufficiency or overload is detrimental to cellular function [30, 33]. Herein, we demonstrate that not only is mitochondrial calcium homeostasis critical for survival of LSCs, but that the steady-state requirement for calcium is substantially higher in venetoclax-resistant LSCs. Intriguingly, this observation may provide a unique opportunity to develop improved clinical strategies for patients who are refractory or relapse following venetoclax-based therapy.

## Results

### Venetoclax Responsiveness is Associated with Intrinsic Differences in Calcium Pathway Signaling

We first validated previous literature that has established the wide range of non-canonical functions of BCL-2. Through an unbiased proteomics screen using BIO-ID, we confirmed the diversity of BCL-2 functions which included proximity interactors at various cellular locations including the ER, mitochondria, nucleus and golgi apparatus (Supplementary Fig. 1A). Further, analysis of proteins detected by BIO-ID showed strong enrichment for pathways involved in calcium transport (Fig. 1A). Next, we reasoned that if calcium signaling is relevant to the action of venetoclax, then calcium biology properties may be different in venetoclax sensitive versus resistant LSCs. To investigate this issue, LSCs were enriched from primary human AML specimens as previously described using labeling of reactive oxygen species (ROS) followed by flow cytometric sorting to isolate ROS-low cells [8, 34]. Venetoclax sensitivity was defined as previously described [5] and validated using viability assays (Supplementary Fig. 2A). Initial comparisons were performed using bulk RNA-seq (Supplementary Fig. 1B-E). The data show significant enrichment of multiple calcium-mediated signaling genesets in venetoclax resistant LSCs, where the “GO-BP Calcium Mediated Signaling Pathway” was the most prevalent (Fig. 1B, Supplementary Fig. 1E) [5].

**Figure 1.**
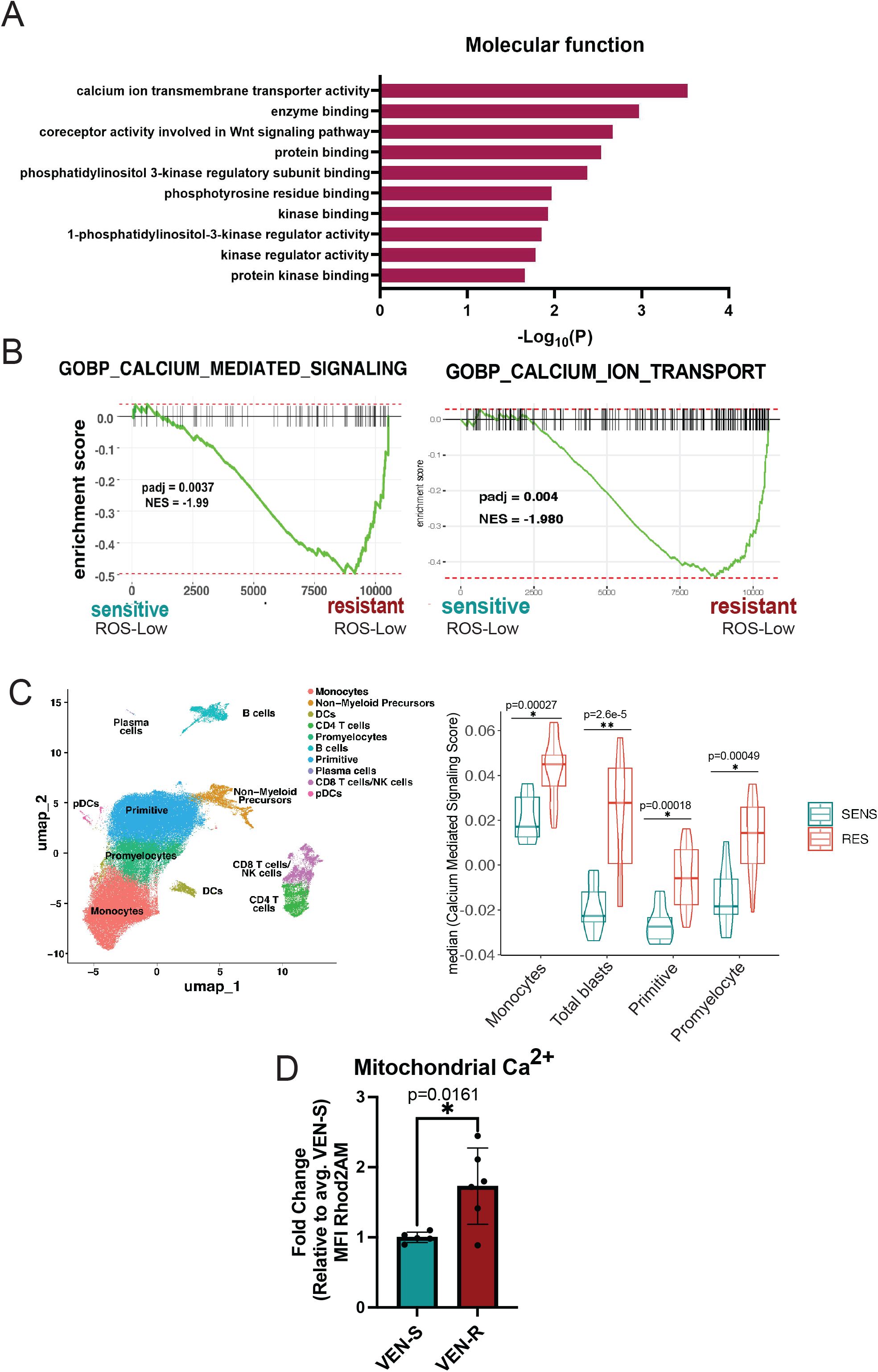
Venetoclax Responsiveness is Associated with Intrinsic Differences in Calcium Pathway Signaling. **A.** BCL-2 interacts with signaling proteins using BioID in T-REx HEK293 cells. Pathway enrichment analysis of BCL-2 proximity interactors involved in cellular signaling. Gene ontology (Molecular Function) analysis shows the top 10 enrichment pathways. **B.** Enrichment plot for GOBP Calcium Mediated Signaling pathway (198 genes) and GOBP Calcium Ion Transport (455 genes) in N=7 venetoclax sensitive primary human AML ROS-Low cells and N=5 venetoclax resistant primary human AML ROS-Low cells **C.** Projection of all samples with cluster assignments annotated. In various populations of tumor including, monocytes, primitive cells, promyelocytes, and total blasts (a combination of monocytes, primitive cells and promyelocytes) GO-BP Calcium Mediated Signaling Pathway expression was analyzed using Seurat’s AddModuleScore function in sensitive (n=8) versus resistant (n=12) AML patient specimens with median plotted. Significance was determined using t-test. **D.** Mitochondrial calcium levels in venetoclax resistant primary human AML ROS-low cells compared to sensitive ROS-low cells presented as mean fluorescence intensity (MFI). N=5 venetoclax sensitive (AML 2,4,6,7,18) and n=6 venetoclax resistant (AML 1,3,11,12,13,14) cells. Data are presented as mean +/-SD. Significance was determined using two-tailed unpaired t-test.

Given the significant heterogeneity observed within primary AML tumors, we next examined how calcium signaling may differ within varying subpopulations using single-cell transcriptomic analyses. The method employed, CITE-Seq (cellular indexing of transcriptomes and epitopes), permits global transcriptomic analyses coupled with high-resolution immunophenotyping. Comparison of venetoclax sensitive (n=8), and resistant (n=14) AML specimens is shown in Fig. 1C, and Supplementary Fig. 1F-J. As expected from bulk RNA-seq findings (Fig. 1B), the “GO-BP Calcium Mediated Signaling” pathway was enriched in venetoclax-resistant specimens (Fig. 1C, Supplementary Fig. 1J). Interestingly, this pathway was enriched in multiple tumor subpopulations including the primitive, monocytic, promyelocytic and the total blast compartment (a compilation of primitive, monocytic, and promyelocytic), suggesting changes in intracellular calcium signaling are a consistent feature of venetoclax resistant cells.

For our subsequent studies, we primarily focused on mitochondrial calcium given its critical role in regulating cellular metabolism and OXPHOS activity. To examine mitochondrial calcium, Rhod2AM labeling was used on ROS-low cells from sensitive (n=5) and resistant (n=6) AML specimens. As shown in Fig. 1D, levels of mitochondrial calcium were significantly higher in venetoclax-resistant specimens. We confirmed mitochondrial localization of the Rhod2AM dye using confocal microscopy which showed co-localization of the dye with mitochondria (Supplementary Fig. 1K). These data are consistent with our previous empirical observations, showing that venetoclax-resistant LSCs have substantially increased mitochondrial metabolism, with more fatty acid oxidation-driven TCA cycle activity, and elevated OXPHOS activity as mitochondrial calcium is required for these processes [5, 9]. Taken together the data suggest that venetoclax sensitive versus resistant LSCs have intrinsically different calcium requirements. The data also imply that mitochondrial calcium levels may be functionally linked to venetoclax resistance in LSCs.

### BCL-2 Inhibition Causes Mitochondrial Calcium Changes Associated with SERCA Disruption in Venetoclax Sensitive LSCs

Given the apparent differences in calcium biology between venetoclax sensitive versus resistant AML specimens, we next investigated the role of venetoclax in modulating calcium signaling in responsive LSCs. ROS-low cells were isolated from venetoclax sensitive specimens, treated with venetoclax, and analyzed by Rhod2AM labeling. Notably, 16-hour treatment with venetoclax led to significant changes in viability (Supplementary Fig. 2A). Therefore, shorter time points (3 hours of treatment) were subsequently employed where no significant changes in mitochondrial membrane potential or viability were evident (Supplementary Fig. 2B, 2C). This approach abrogates the potential confounding effect of overt cell death on the results. Venetoclax significantly increased mitochondrial calcium levels in sensitive primary human AML LSCs (Fig. 2A). Importantly, these changes were not observed in the ROS-high population indicating that venetoclax-induced calcium flux is relatively specific to the LSC compartment (Supplementary Fig. 2D). To confirm whether the venetoclax mediated change in mitochondrial calcium was an on-target effect of BCL-2 inhibition, we utilized electroporation of siRNA to knock-down BCL-2 (Supplementary Fig. 2E). Notably, inhibition of BCL-2 in sensitive LSCs phenocopied the effects of venetoclax on mitochondrial calcium levels (Fig. 2B). Further, calcium flux into the mitochondria preceded the onset of overt apoptosis or cell death (Supplementary Fig. 2F-G). To determine if changes in mitochondrial calcium could contribute to venetoclax-mediated inhibition of OXPHOS activity, we performed seahorse assays at the same time point. As shown in Supplementary Fig. 2H-I, both venetoclax treatment and genetic inhibition of BCL-2 decreased OXPHOS activity in sensitive LSCs. Taken together, these data suggest that mitochondrial calcium overload arising from BCL-2 inhibition may contribute to OXPHOS inhibition.

**Figure 2.**
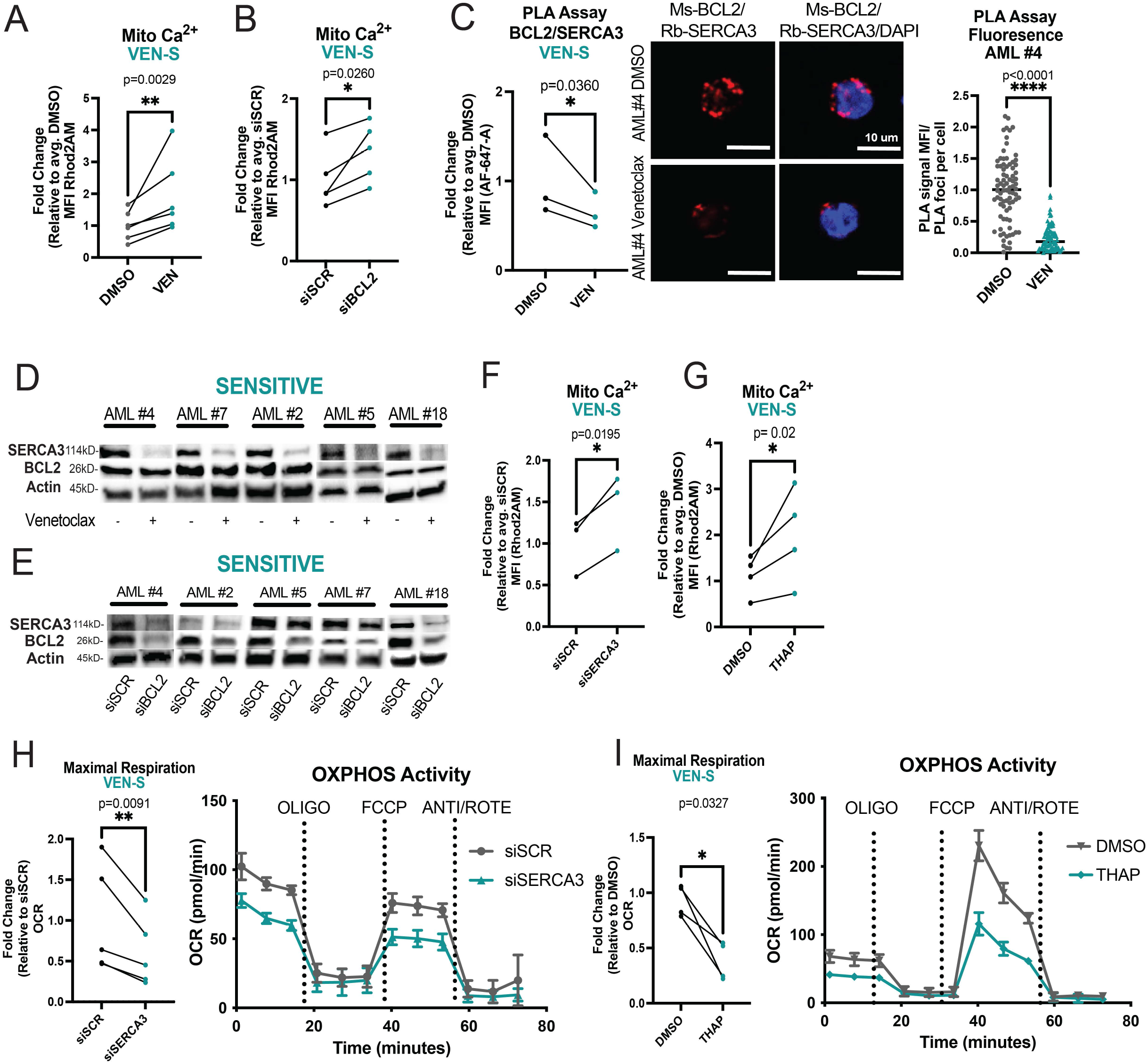
BCL-2 Inhibition Causes Mitochondrial Calcium Changes Associated with SERCA Disruption in Venetoclax Sensitive LSCs. **A.** Mitochondrial calcium content presented as mean fluorescence intensity (MFI) after venetoclax treatment (500nM, 3 hours). N=6 venetoclax sensitive primary human AML ROS-low cells (AML#2,4-7,18). Significance was determined using two-tailed ratio paired t-test. **B.** Primary human ROS-low AML cells analyzed 36 hours post electroporation with indicated siRNA. Mitochondrial calcium after genetic inhibition of BCL-2. N=5 venetoclax sensitive specimens (AML# 2,4-7). Significance was determined using two-tailed ratio paired t-test. **C.** BCL-2 and SERCA3 proximity interaction in primary human ROS-low AML cells after venetoclax treatment (500nM, 3 hours) as measured by PLA assay with flow cytometry (n=3, AML #2,4,7). Significance was determined using two-tailed ratio paired t-test. Confocal microscopy to confirm PLA assay results. Representative images from AML #4. Red signal is positive signal from PLA assay (FarRed) while blue signal is from Dapi staining. Signal was quantified using ImageJ for MFI of loci. N=82 loci for DMSO and N=77 loci for venetoclax. Data are presented as mean with individual data points. Significance was determined using two-tailed unpaired t-test. **D.** Western blot of SERCA3 and BCL-2 levels upon venetoclax treatment (500nM, 3 hours) in venetoclax sensitive primary human AML ROS low cells. Anti-actin antibody was used as loading control, anti-BCL-2 and anti-SERCA3 antibodies were used to determine protein levels of BCL-2 and SERCA3 respectively. Quantification presented in supplementary figure 2J **E.** Western blot of SERCA3 levels upon BCL-2 inhibition via siRNA electroporation in primary human ROS-low AML cells 36 hours post infection. Anti-actin antibody was used as loading control, anti-BCL-2 and anti-SERCA3 antibodies were used to determine protein levels of BCL-2 and SERCA3 respectively. Quantification presented in supplementary figure 2K **F.** Mitochondrial calcium 36 hours post siRNA electroporation of primary human ROS-low AML cells. N=3 (AML#2,5,6). Significance determined using two-tailed ratio paired t-test. **G.** Mitochondrial calcium content after thapsigargin treatment (500nM, 3 hours). N=4 primary human ROS-low AML cells (AML# 2,5-7). Significance determined using two-tailed ratio paired t-test. **H.** OCR after genetic knockdown of SERCA3 36 hours post infection in venetoclax sensitive primary human ROS-low AML cells. N=5 (AML# 2,4-7). Significance was determined using two-tailed ratio paired t-test. **I.** OCR after thapsigargin treatment (500nM, 3 hours). N=4 (AML# 2,5-7) in venetoclax sensitive primary human ROS-low AML cells. Significance was determined using two-tailed ratio paired t-test.

Notably, previous studies in other cell types (DLBCL and pancreatic acinar cell lines) showed that in contrast to our findings venetoclax does not perturb intracellular calcium signaling [35, 36]. This has been ascribed to the fact that venetoclax is a BH3 mimetic which primarily disrupts the BH3 domain, while a majority of BCL-2 binding interactions with intracellular calcium channels are thought to occur at the BH4 domain. Specifically, the key ER/mitochondrial calcium channels VDAC (responsible for influx of calcium and other metabolites into mitochondria) [37], and IP3R (responsible for calcium efflux out of the ER) [18] have been shown to lose binding with BCL-2 upon perturbation of the BH4 domain. In contrast, the binding relationship between SERCA3, the main protein responsible for calcium influx at the ER, and the BH domains of BCL2 has not been fully elucidated [16]. Thus, we investigated whether venetoclax could perturb intracellular calcium signaling by modulating SERCA3 activity. We utilized proximity ligation assays (PLA) to first establish the relationship between the main calcium regulatory proteins and BCL-2. As expected, BCL-2 showed proximity interactions with SERCA3, VDAC and IP3R in venetoclax sensitive ROS-low cells (Supplementary Fig. 2L-N). Additionally, as a control for non-specific antibody binding, PLAs were performed using primary AML cells with IgG controls and BCL-2 knockdown, which confirmed the specificity of the assay (Supplementary Fig. 2L-M). Notably, the interaction with SERCA3 was the most consistent and statistically significant interaction among multiple samples (Supplementary Fig. 2N). Further, venetoclax treatment consistently and significantly decreased the proximity interaction of BCL2 and SERCA3 in sensitive LSCs (Fig. 2C, Supplementary Fig. 2O) as measured by both confocal microscopy and flow cytometry-based PLA.

To determine how venetoclax may perturb BCL-2/SERCA3 interaction in sensitive LSCs, we first used immunoblotting to assess whether protein levels of BCL-2 or SERCA3 were changed upon treatment. We observed a strong reduction in SERCA3 protein levels after 3 hours of venetoclax treatment, suggesting that the loss of PLA signal was due to SERCA3 degradation and/or reduced expression (Fig. 2D, Supplementary Fig. 2J). siRNA-mediated knockdown of BCL-2 also led to decreased SERCA3 levels, supporting an on-target role of venetoclax in regulating SERCA3 (Fig. 2E, Supplementary Fig. 2K). As venetoclax treatment did not lead to significant changes in ATP2A3 (the gene encoding SERCA3) transcript levels (Supplementary Fig. 2Q), we hypothesized proteolysis may be the mechanism by which venetoclax modulates SERCA3 degradation in sensitive LSCs. Preincubation with MG-132, a proteasome inhibitor, rescued SERCA3 levels in two of five samples tested indicating proteolysis may contribute to venetoclax-mediated SERCA3 loss in at least some AML specimens (Extended Data Fig. 2T).

To test whether loss of SERCA3 is functionally relevant for venetoclax sensitive LSCs, we employed genetic and pharmacologic inhibition of SERCA. Inhibition of SERCA3 by siRNA-mediated knockdown (Supplementary Fig. 2R-S) or treatment with the SERCA inhibitor thapsigargin led to increased mitochondrial calcium (Fig. 2F, 2G) and decreased OXPHOS activity (Fig. 2H, 2I) in venetoclax sensitive ROS-low AML cells, thereby phenocopying the effects of BCL-2 inhibition. Additionally, at the early time points assessed, there were no changes in overt cell death upon genetic or pharmacologic SERCA inhibition (Supplementary Fig. 2P). To assess the effect of SERCA inhibition on LSC activity, we utilized patient derived xenograft (PDX) models to functionally measure LSC activity. Briefly, venetoclax sensitive primary AML specimens were treated overnight with vehicle or thapsigargin, then transplanted into immunodeficient mice (Supplementary Fig. 2U). As shown in Figure 2U, we observed reduction in LSC engraftment potential of thapsigargin treated cells. Notably, thapsigargin did not affect engraftment of normal mobilized peripheral blood samples in NSG-S mice, suggesting SERCA inhibition is a vulnerability in venetoclax sensitive LSCs (Supplementary Fig. 2V). Taken together, these data indicate that SERCA inhibition is able to phenocopy the effects of BCL-2 inhibition on mitochondrial calcium, OXPHOS activity and LSC function in venetoclax sensitive AML specimens.

### Reducing Mitochondrial Calcium Levels Targets Venetoclax Resistant LSCs

As the data in Figure 2 indicate a link between venetoclax activity and calcium signaling, we next examined venetoclax-resistant specimens. In contrast to our findings with drug sensitive specimens, venetoclax treatment did not induce any changes in mitochondrial calcium, BCL-2/SERCA3 proximity interactions, or SERCA3 levels in resistant LSCs (Fig. 3A-C, Supplementary Fig. 3A-B). In addition, transcriptomic analyses showed that SERCA3 (ATP2A3) expression was decreased in resistant LSCs whereas expression of MCU and MCUB, channels responsible for importing calcium into the mitochondrial matrix, were increased (Fig. 3D, Supplemental Fig. 3C) These data support the concept that ven-resistant LSCs exist in a more active metabolic state in which intrinsically higher levels of mitochondrial calcium are required and therefore more calcium is sequestered to the mitochondria.

**Figure 3.**
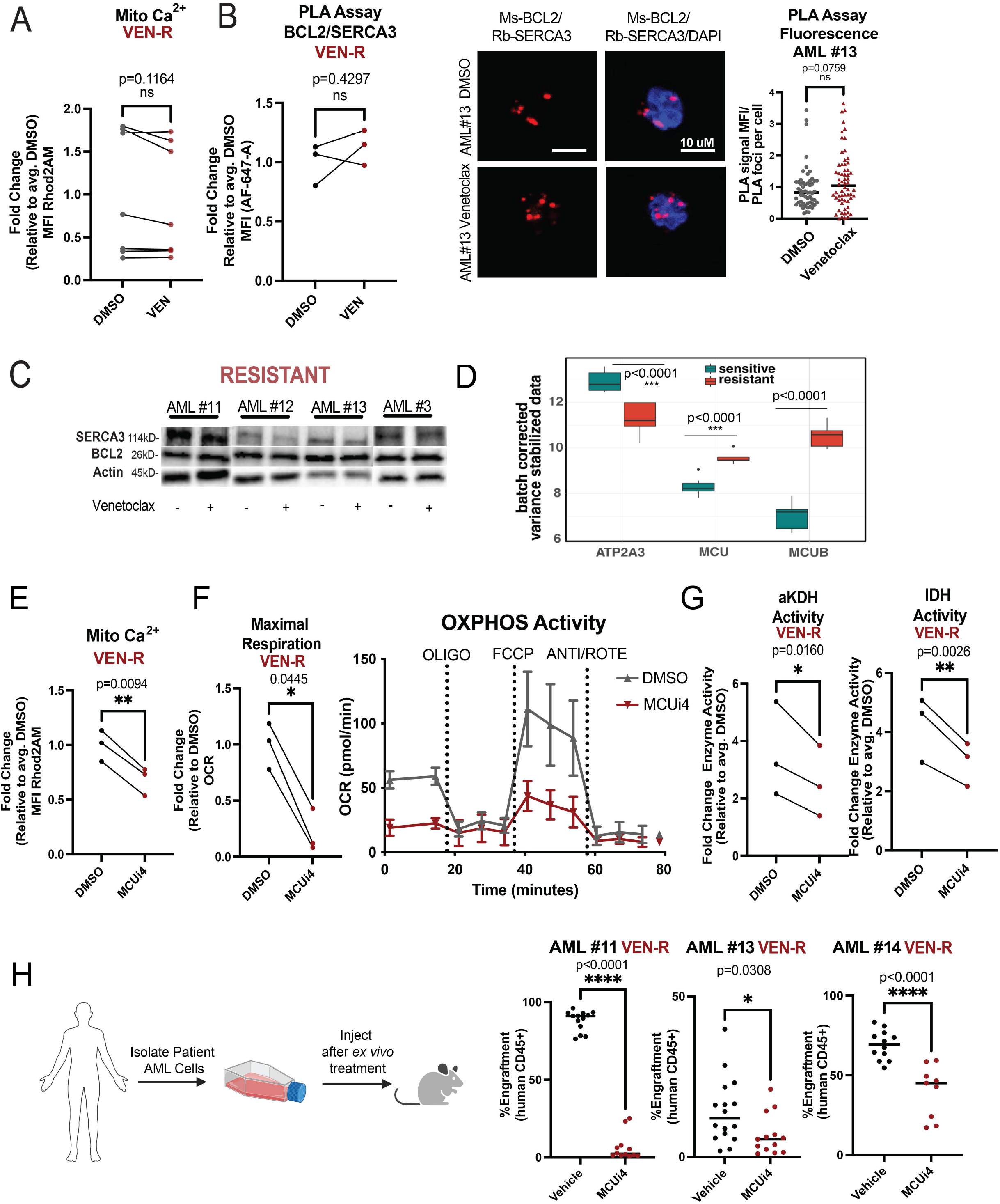
Reducing Mitochondrial Calcium Levels Targets Venetoclax Resistant LSCs. **A.** Mitochondrial calcium content presented as mean fluorescence intensity (MFI) after venetoclax treatment (500nM, 3 hours). N=7 venetoclax resistant primary human AML ROS-low cells (AML# 11-14, 19-21). Significance was determined using two-tailed ratio paired t-test. **B.** BCL-2 and SERCA3 proximity interaction in venetoclax resistant primary human AML ROS-low cells after venetoclax treatment (500nM, 3 hours) as measured by PLA assay with flow cytometry (n=3, AML #11-13). Significance was determined using two-tailed ratio paired t-test. Confocal microscopy to confirm PLA assay results. Representative images from AML #13. Red signal is positive signal from PLA assay (FarRed) while blue signal is from Dapi staining. Signal was quantified using ImageJ for MFI of loci. N=56 foci for DMSO and N=62 foci for venetoclax. Data are presented as mean with individual data points. Significance was determined using two-tailed unpaired t-test. **C.** Western blot of SERCA3 and BCL-2 levels upon venetoclax treatment (500nM, 3 hours) in venetoclax resistant primary human AML ROS-Low cells. Anti-actin antibody was used as loading control, anti-BCL-2 and anti-SERCA3 antibodies were used to determine protein levels of BCL-2 and SERCA3 respectively. Quantification presented in supplementary figure 3B **D.** Boxplot of specific genes of interest. All genes shown were significantly different between resistant and sensitive samples with p-values shown on figure. N=7 venetoclax sensitive ROS-Low cells and N=5 venetoclax resistant ROS-Low cells. **E-G.** Venetoclax resistant primary human AML ROS-Low cells (n=3, AML#11-13) treated with MCUi4 for 16 hours at 5uM. **E.** Mitochondrial calcium content after MCUi4 treatment presented as mean fluorescence intensity (MFI). Significance was determined using two-tailed ratio paired t-test. **F.** OCR after MCUi4 treatment. Significance was determined using two-tailed ratio paired t-test. **G.** Isocitrate dehydrogenase activity and alpha keto-glutarate dehydrogenase activity after MCUi4 treatment. Significance was determined using two-tailed ratio paired t-test. **H.** MCUi4 treatment (5uM, 16 hours, ex vivo) and subsequent engraftment potential of venetoclax resistant primary human AML specimens after transplantation into immune-deficient mice. N=13,16,12 for vehicle control group and n=12,12,9 for MCUi4 treatment group for AML 11,13 and 14 respectively. 2 million cells injected per mouse and engraftment was assessed between 4-8 weeks. Data are presented as mean with individual data points. Significance was measured by two-tailed unpaired t-test.

Given the increased expression of MCU, higher basal mitochondrial metabolism and higher mitochondrial calcium levels in resistant LSCs, we hypothesized that MCU inhibition could be a strategy to reduce mitochondrial calcium and thereby target venetoclax-resistant LSCs. To test this concept, we first employed laboratory grade pharmacological inhibitors of MCU (MCUi4, Ru265) and siRNA mediated genetic inhibition in venetoclax resistant LSCs. Treatment with MCUi4, a compound that inhibits MICU1, a protein that controls the activity of MCU [38] led to decreased mitochondrial calcium (Fig. 3E), decreased OXPHOS activity (Fig. 3F) and decreased activity of the calcium-dependent metabolic dehydrogenases alpha-ketoglutarate dehydrogenase and isocitrate dehydrogenase (Fig. 3G). Additionally, Ru265, a compound that directly binds to the matrix side of MCU and inhibits its activity [39] showed similar effects (Supplementary Fig. 3H-J). Consistent with the pharmacological data, siRNA mediated knockdown of MCU showed decreased OXPHOS activity (Supplementary Fig. 3L-N). To determine whether MCU inhibition could directly affect venetoclax-resistant LSCs, we performed patient-derived xenograft (PDX) assays using venetoclax-resistant primary AML specimens. As shown in Figure 3H, we observed a consistent reduction in LSC engraftment potential of MCUi4 treated cells. These data were corroborated by Ru265 treatment which led to decreased colony forming abilities in venetoclax resistant specimens (Supplementary Fig. 3K). Importantly, at the time points assessed in the above experiments, there were no changes in viability upon treatment indicating the effects observed from MCU inhibition were before overt cell death (Supplementary Fig. 3D-E). Further, both MCUi4 and Ru265 did not affect engraftment or colony-forming abilities of normal mobilized peripheral blood samples, suggesting MCU inhibition is a specific vulnerability in venetoclax resistant LSCs (Supplementary Fig. 3F-G).

### Mitoxantrone Inhibits Mitochondrial Metabolism and Colony Formation in Venetoclax Resistant LSCs

To determine whether inhibition of mitochondrial calcium uptake could be clinically applicable, we examined the literature for FDA approved mitochondrial calcium inhibitors. Several agents have been shown to inhibit mitochondrial calcium uptake including doxycycline/minocycline [40], benzothenium chloride [41] and mitoxantrone [42]. Of these agents, only the well charactered chemotherapy drug mitoxantrone (mitox) is known to be a direct MCU inhibitor. While mitox is primarily known as a DNA-damaging agent, a recent study reported a single amino acid substitution in MCU rendered mitoxantrone unable to bind and inhibit mitochondrial calcium uptake [42]. We therefore investigated mitox as an agent to suppress calcium transport in venetoclax-resistant LSCs. As shown in Fig. 4A-C, mitox treatment of venetoclax-resistant specimens demonstrated a consistent reduction in mitochondrial calcium, a marked decrease in OXPHOS activity, and suppression of calcium-dependent dehydrogenases. Importantly, at the relatively low dose employed (100nM) there was no detectable DNA damage observed (as assayed by gamma H2AX labeling, Fig. 4D) or changes in viability (Supplementary Fig. 4A). To determine the functional consequence of mitox treatment, a panel of venetoclax-resistant specimens was tested for colony-forming ability. In all specimens’ doses as low as 10nM completely eradicated colony-forming ability with significant suppression evident at 1nM in three of four specimens (Fig. 4E). Control studies show no significant inhibition of colony-formation for normal stem/progenitor cells derived from mobilized peripheral blood specimens (Fig. 4F).

**Figure 4.**
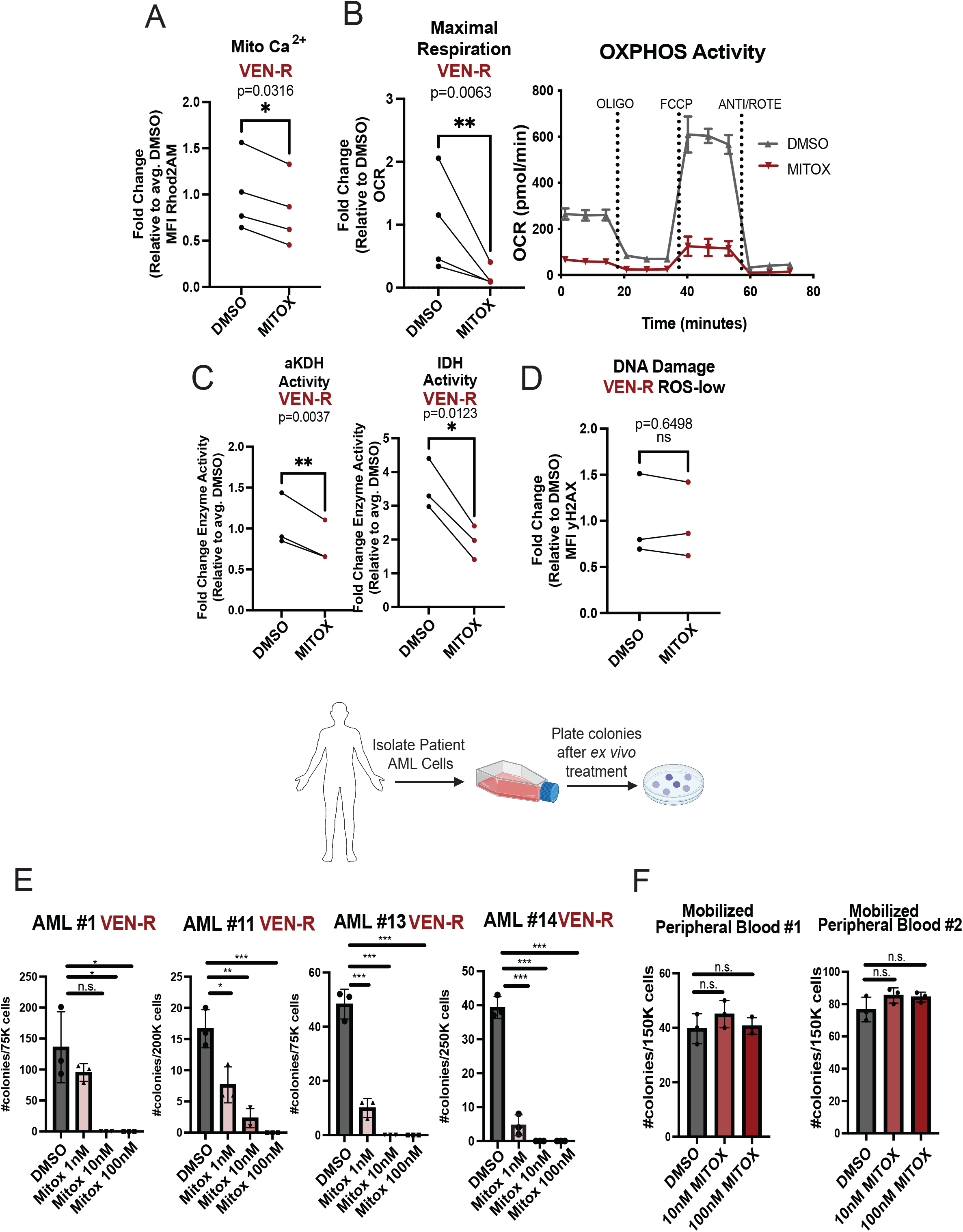
Mitoxantrone Inhibits Mitochondrial Metabolism and Colony Formation in Venetoclax Resistant LSCs. **A-D.** Venetoclax resistant primary human AML ROS-low cells treated with mitoxantrone for 16 hours at 100nM. **A.** Mitochondrial calcium content after mitoxantrone treatment presented as mean fluorescence intensity (MFI). Significance was determined using two-tailed ratio paired t-test. (N=4, AML#11-14) **B.** OCR after mitoxantrone treatment. Significance was determined using two-tailed ratio paired t-test. (N=4, AML#11-14) **C.** Isocitrate dehydrogenase activity and alpha keto-glutarate dehydrogenase activity after mitoxantrone treatment (N=3, AML#11-13). Significance was determined using two-tailed ratio paired t-test. **D.** Gamma H2AX presented as mean fluorescence intensity (MFI). Significance was determined using two-tailed ratio paired t-test. (N=3, AML#11,13,14) **E.** Colony forming units measured by colony formation assays in venetoclax resistant primary human AML specimens after mitoxantrone treatment (1nM,10nM and 100nM, 16 hour treatment, *ex vivo*). Data are presented as mean values +/-SD. Significance was measured by two-tailed unpaired t-test. N=3 per sample per condition. **F**. Colony forming units measured by colony formation assays in mobilized peripheral blood samples after mitoxantrone treatment (DMSO, 10nM or 100nM, 16 hour treatment, *ex vivo*). Data are presented as mean values +/-SD. Significance was measured by two-tailed unpaired t-test. N=3 per sample per condition.

To further investigate the specificity of mitox, we also tested the related anthracycline doxorubicin, and the topoisomerase inhibitor etoposide. Doxorubicin did not inhibit uptake of mitochondrial calcium or OXPHOS activity (Supplementary Fig. 4C-D). Further, dose-response experiments show that while mitox and etoposide were both effective inducers of DNA damage (measured by yH2AX), only mitox was able to reduce mitochondrial calcium in a dose-dependent manner (Supplementary Fig.4E). Functionally, neither doxorubicin nor etoposide were able to substantially suppress the colony forming ability of venetoclax resistant specimens (Supplementary Fig. 4B).

### Mitoxantrone Targets Venetoclax Resistant LSCs

To further investigate the efficacy of mitox, we utilized patient derived xenograft (PDX) models to measure LSC activity. Pre-treatment with mitoxantrone strongly suppressed the engraftment potential of five independent AML specimens (Fig. 5A) that span a variety of mutational backgrounds (KRAS, PTPN11, FLT-3-ITD/TKD, and MLL rearranged, see Table 2). Next, to test in vivo activity of mitox, we transplanted venetoclax-resistant AML specimens into NSG-S immune deficient mice and allowed tumor burden to establish in marrow (∼4 weeks). Animals were then treated with mitox (0.5mg/kg/day) for 4 days followed by sacrifice and assessment of human leukemic cells. This regimen resulted in a modest reduction of primary tumor burden in marrow (Fig. 5B), with no significant weight loss (Supplementary Fig. 5A) or overt toxicity. However, upon secondary transplantation, an assay that directly measures LSC activity, we observed complete suppression of engraftment in leukemic cells derived from mitox treated primary mice (Fig. 5B, Supplementary Fig. 5B). These data indicate effective in vivo targeting of venetoclax-resistant LSCs by mitox.

**Figure 5.**
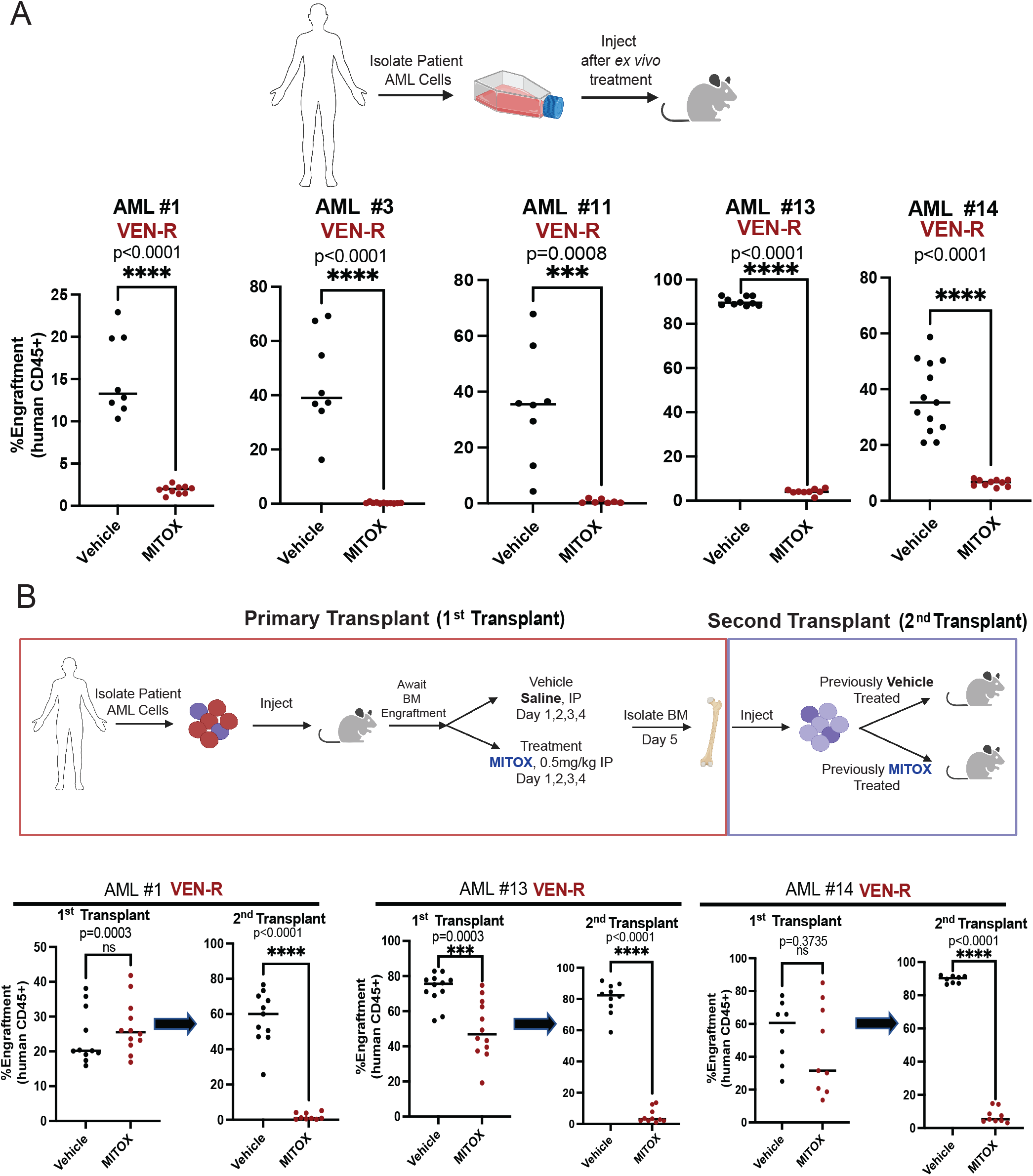
Mitoxantrone Targets Venetoclax Resistant LSCs. **A.** Engraftment potential of venetoclax resistant primary human AML specimens after mitoxantrone treatment (100nM, 16 hours, *ex vivo*) after transplantation into immune-deficient mice. Data are presented as mean values with individual data points. Significance was measured by two-tailed unpaired t-test. n=8,8,8,10,13 for vehicle control group and n=10,11,7,10,10 for mitoxantrone treatment group for AML 1,3,11,13, 14 respectively. **B.** In vivo treatment of NSG-S mice with venetoclax resistant AML tumor burden. Venetoclax resistant specimens were transplanted into immune-deficient mice. Upon at least 20% bone marrow tumor burden, mice were treated with either vehicle (PBS,i.p/day. 4 days) or mitoxantrone (0.5mg/kg/day,i.p. 4 days). On day 5, mice were sacrificed and bone marrow was assessed for tumor burden. Cells were then transplanted again into NSG-S mice for secondary transplants (1million cells per mouse per condition) to assess LSC potential. Data are presented as mean values with individual data points. Significance was measured by two-tailed unpaired t-test. For AML #1, n=11 (vehicle) and n=12 (mitoxantrone) for primary transplants. For AML #1, n=11 (vehicle) and n=9 (mitoxantrone) for secondary transplants. For AML #13, n=12 (vehicle) and n=12 (mitoxantrone) for primary transplants. For AML #13, n=10 (vehicle) and n=10 (mitoxantrone) for secondary transplants. For AML#14, n=8 (vehicle) and n=9 (mitoxantrone) for primary transplants. For AML #11, n=8 (vehicle) and n=9 (mitoxantrone) for secondary transplants.

**Table 1.**
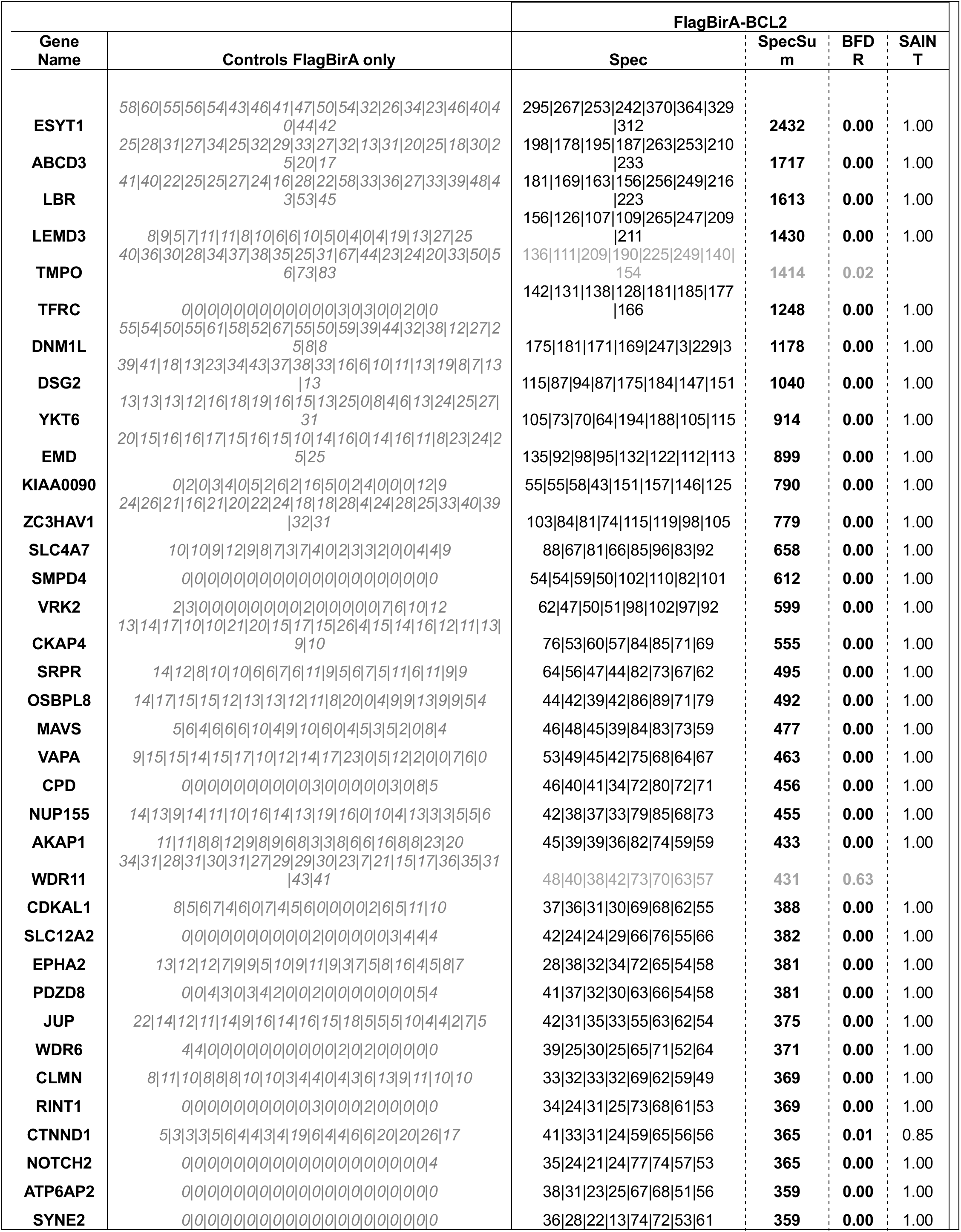
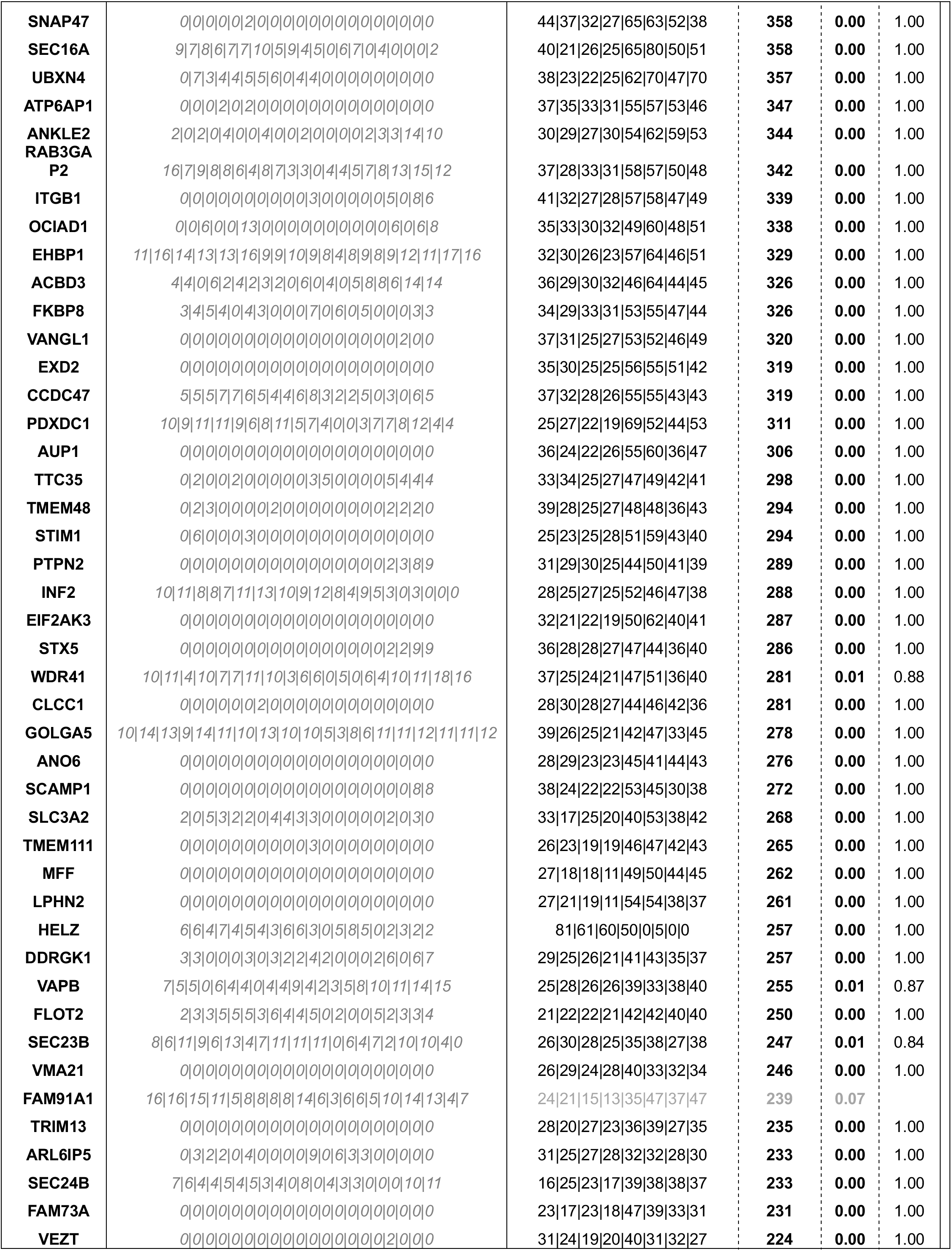
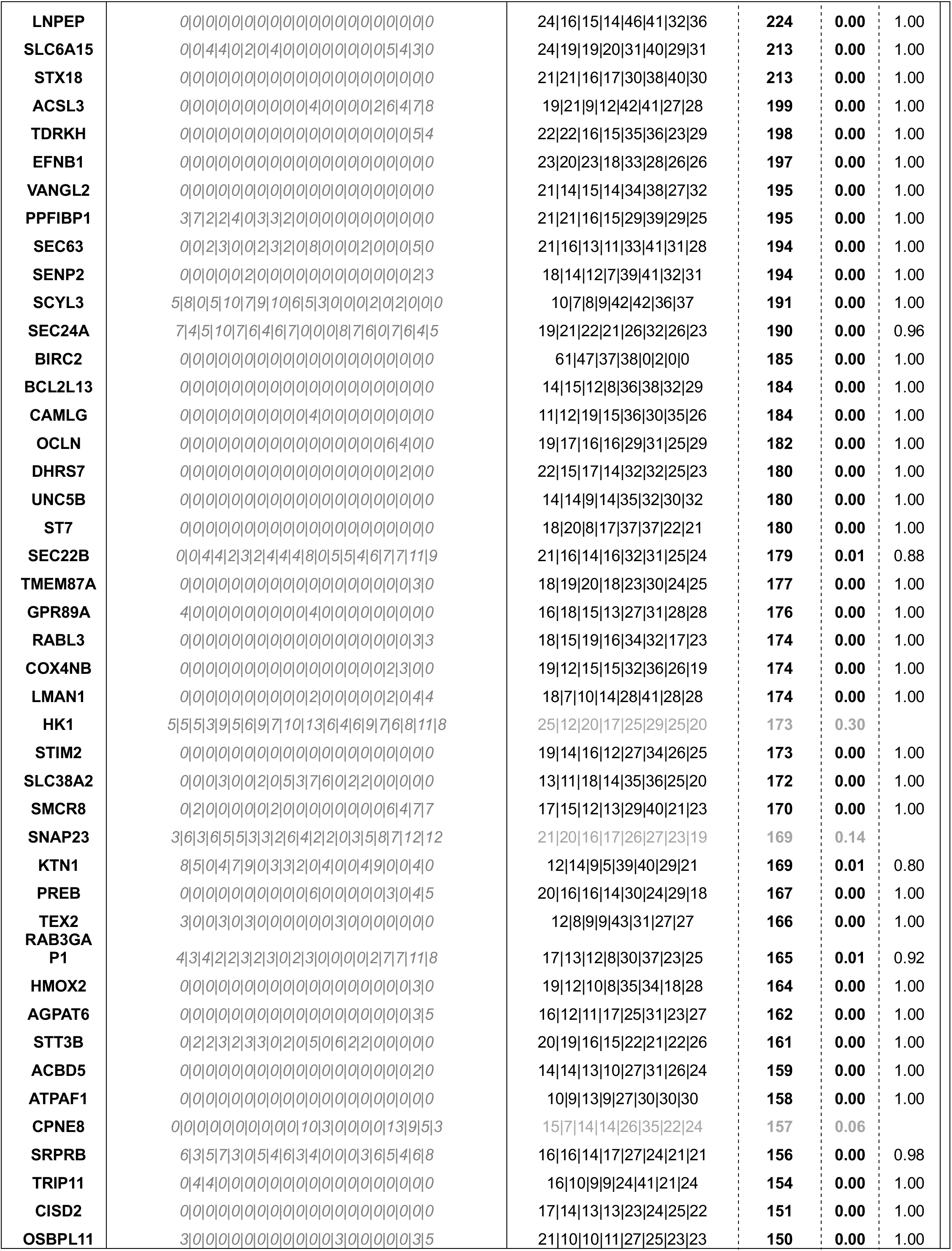
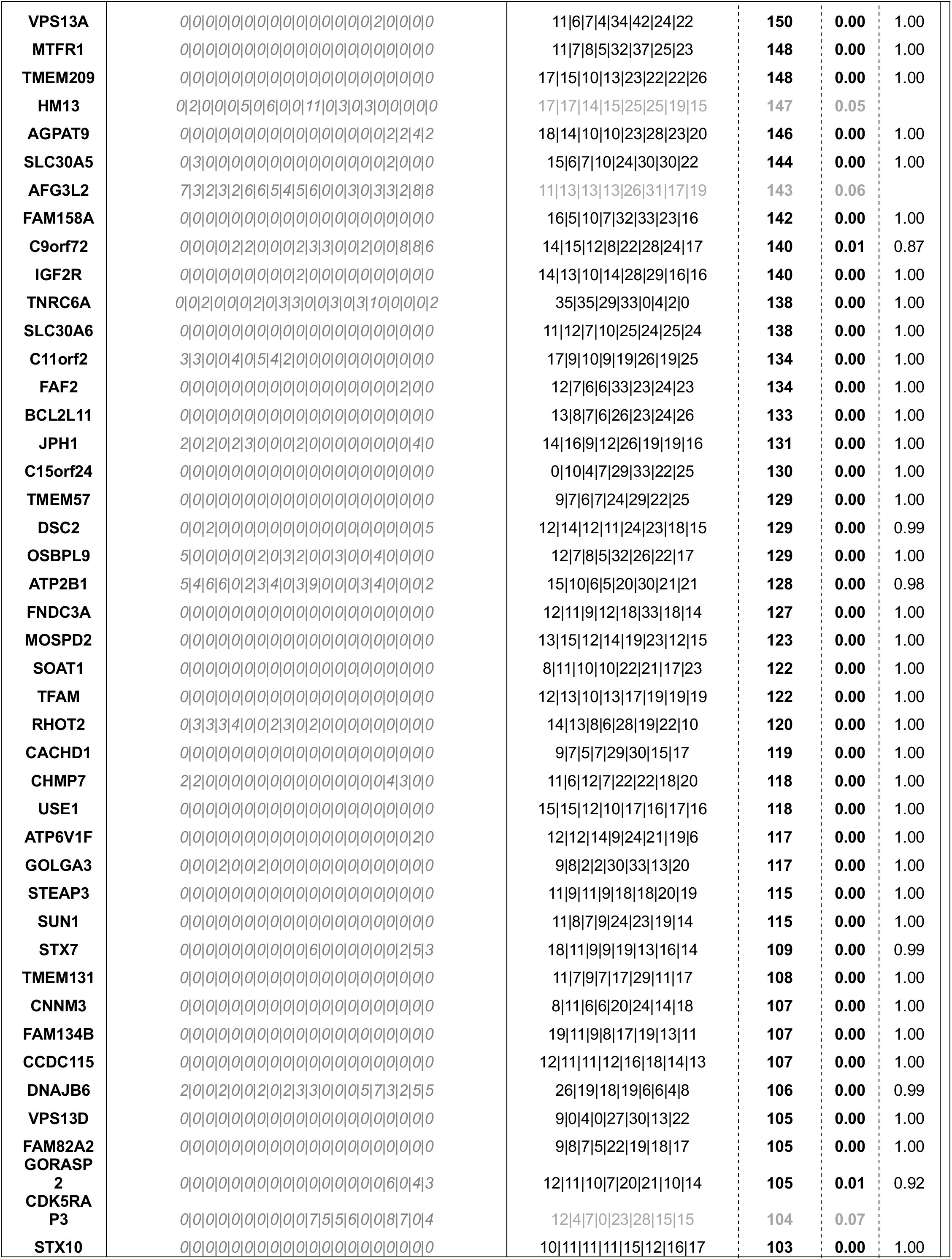
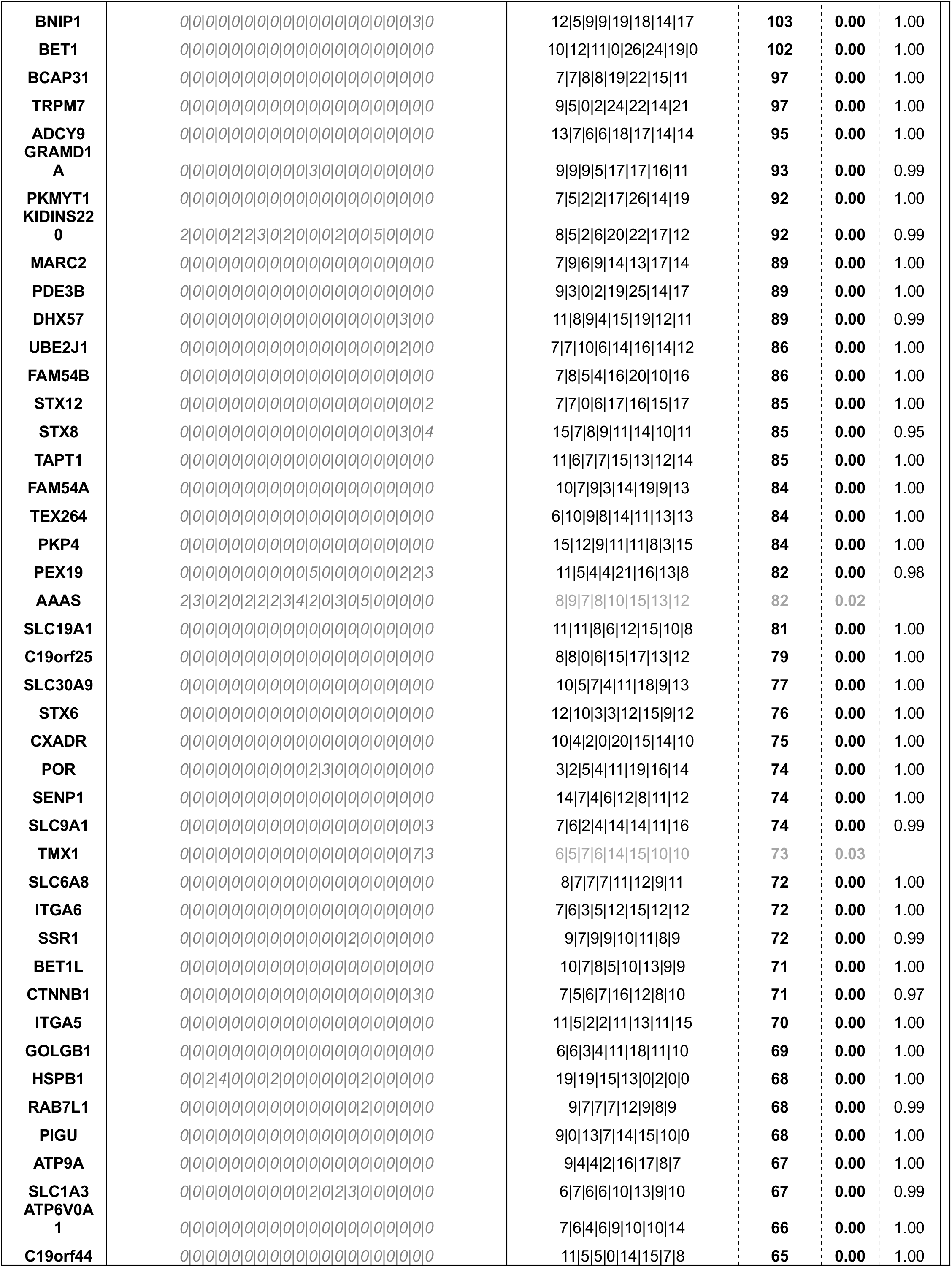
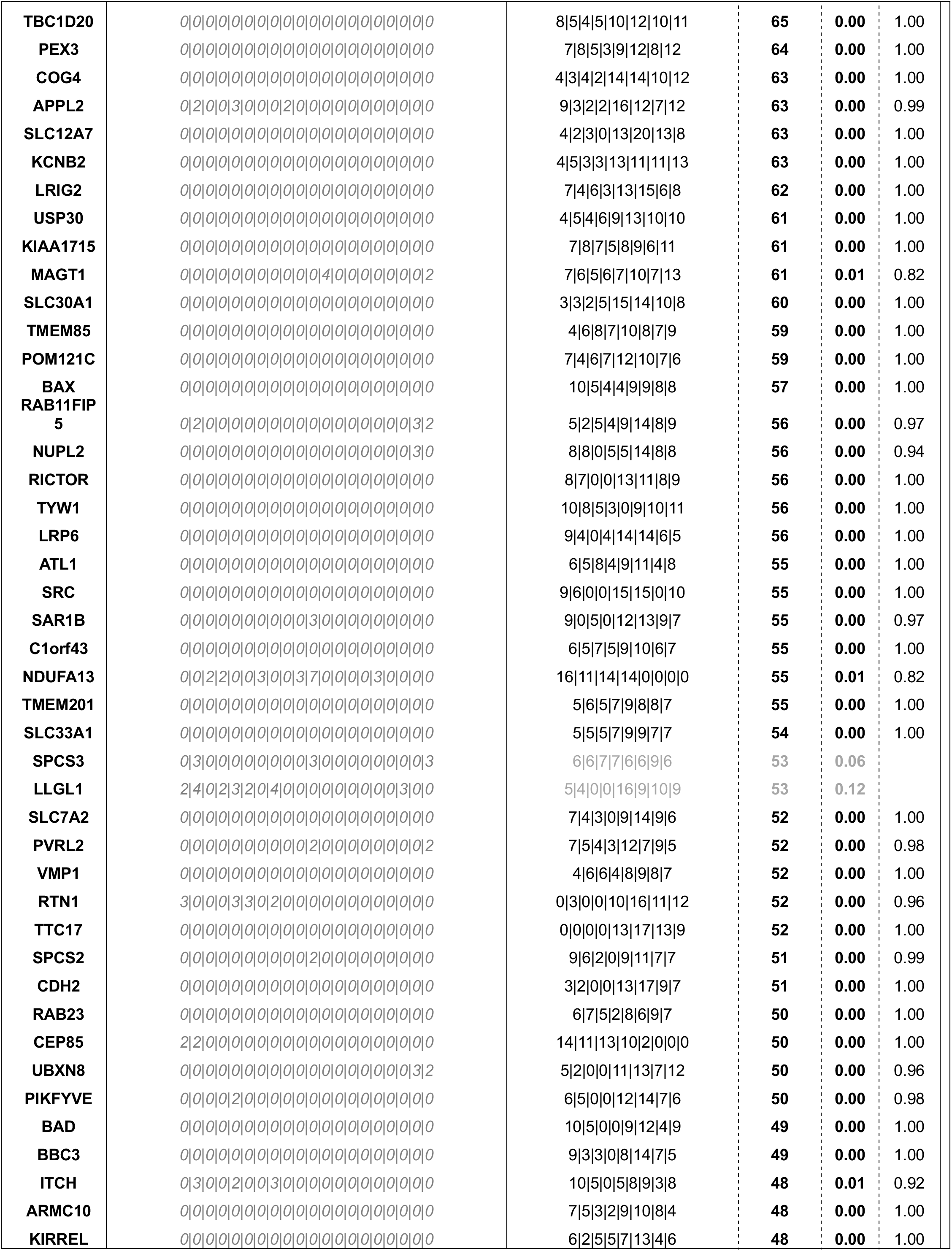
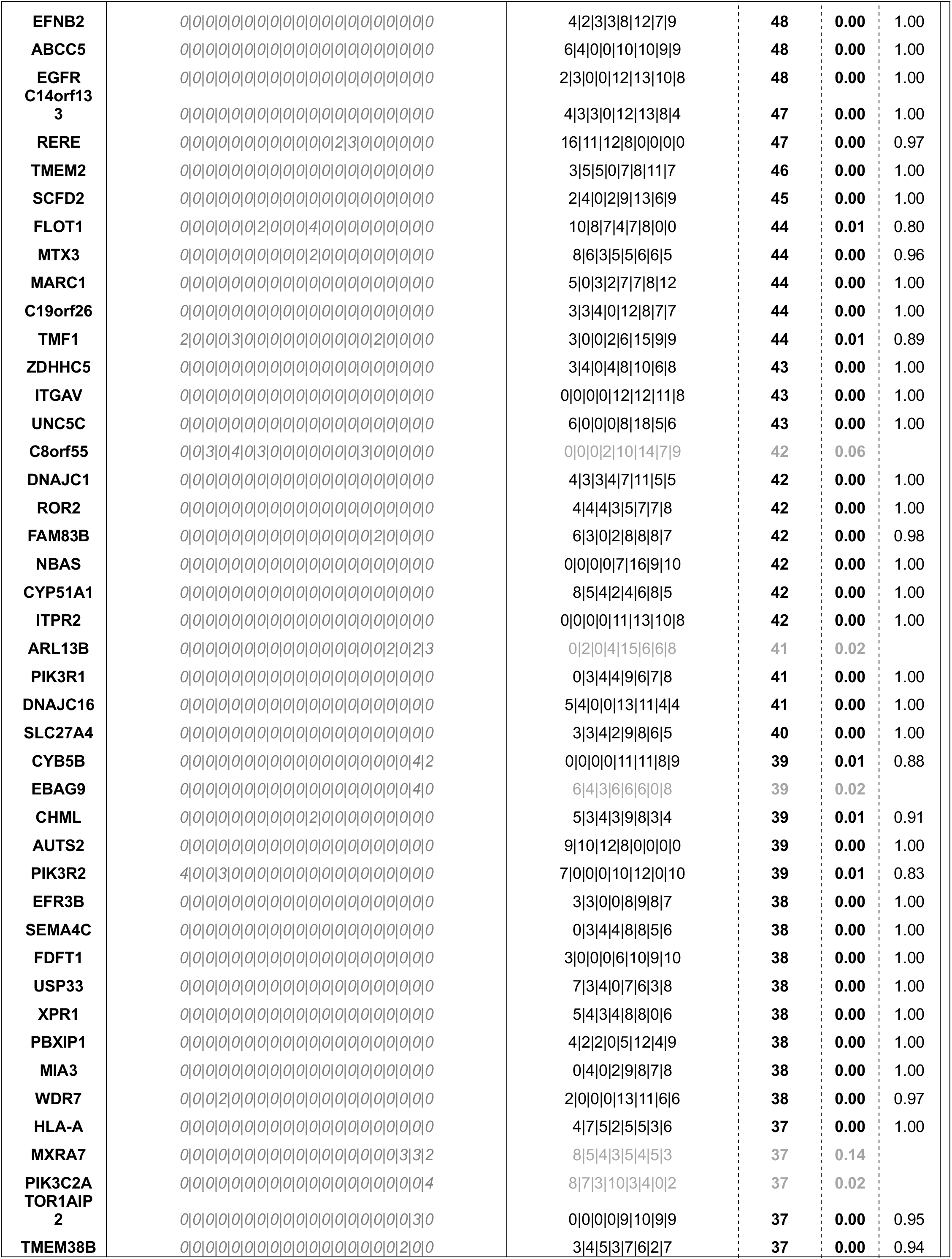
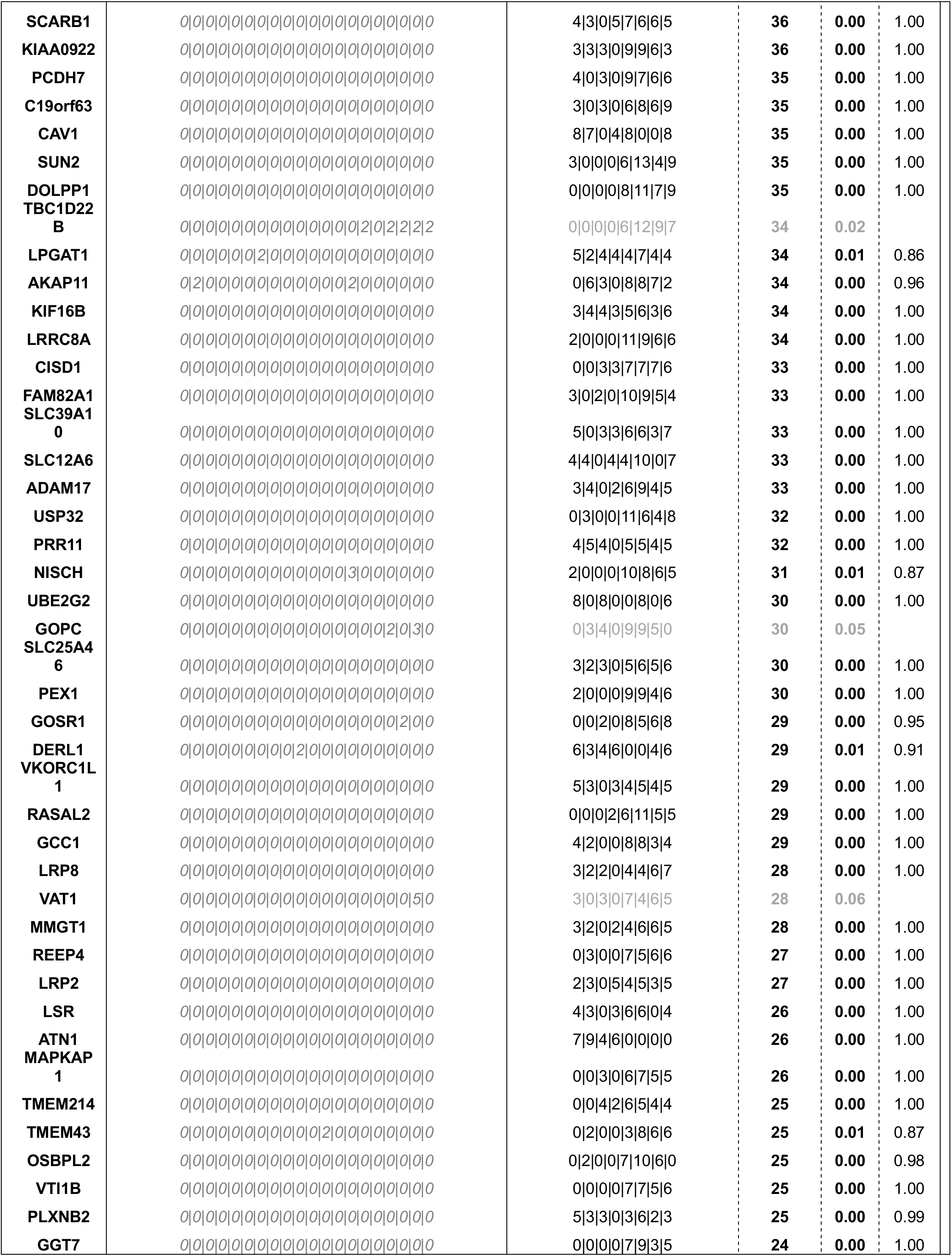
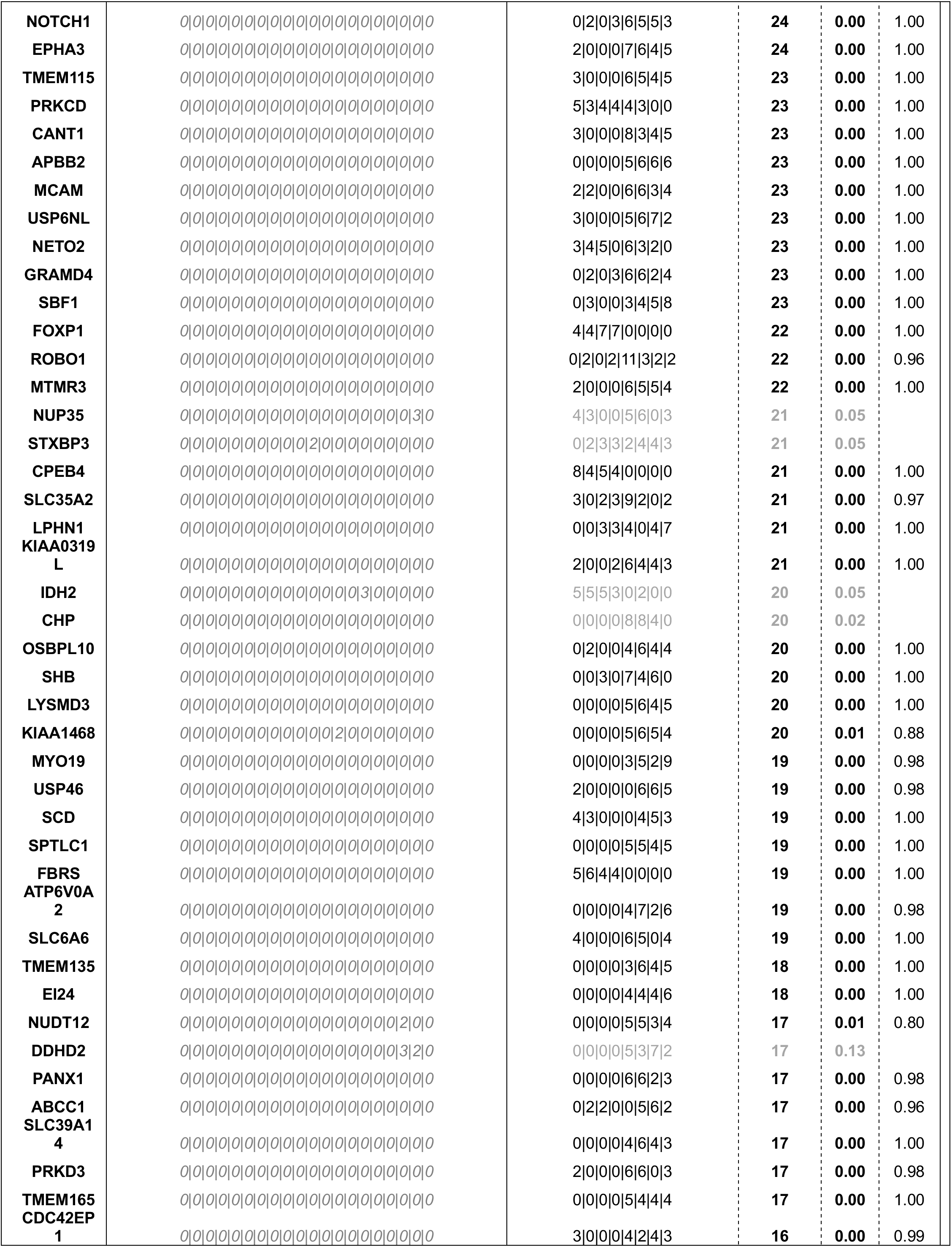
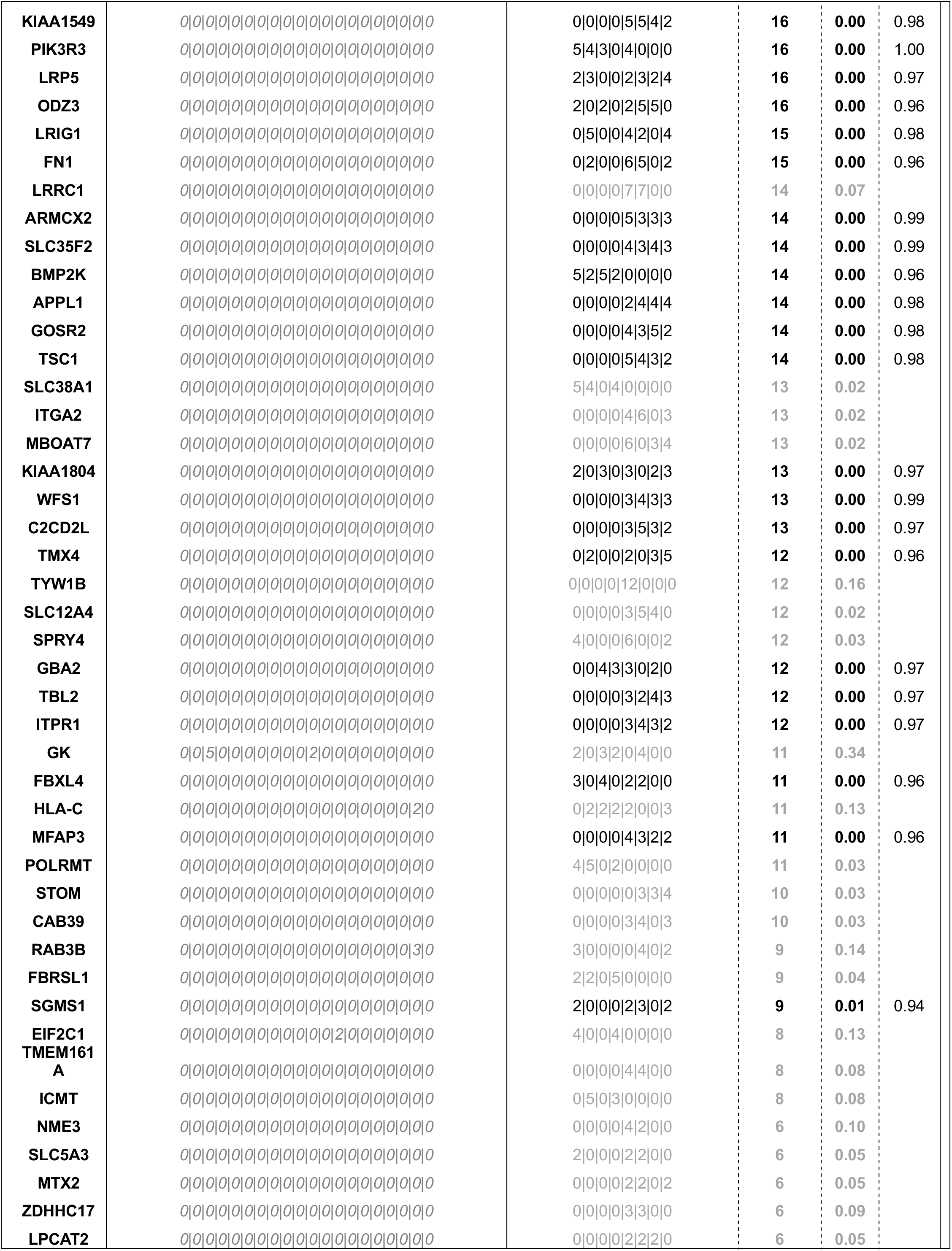
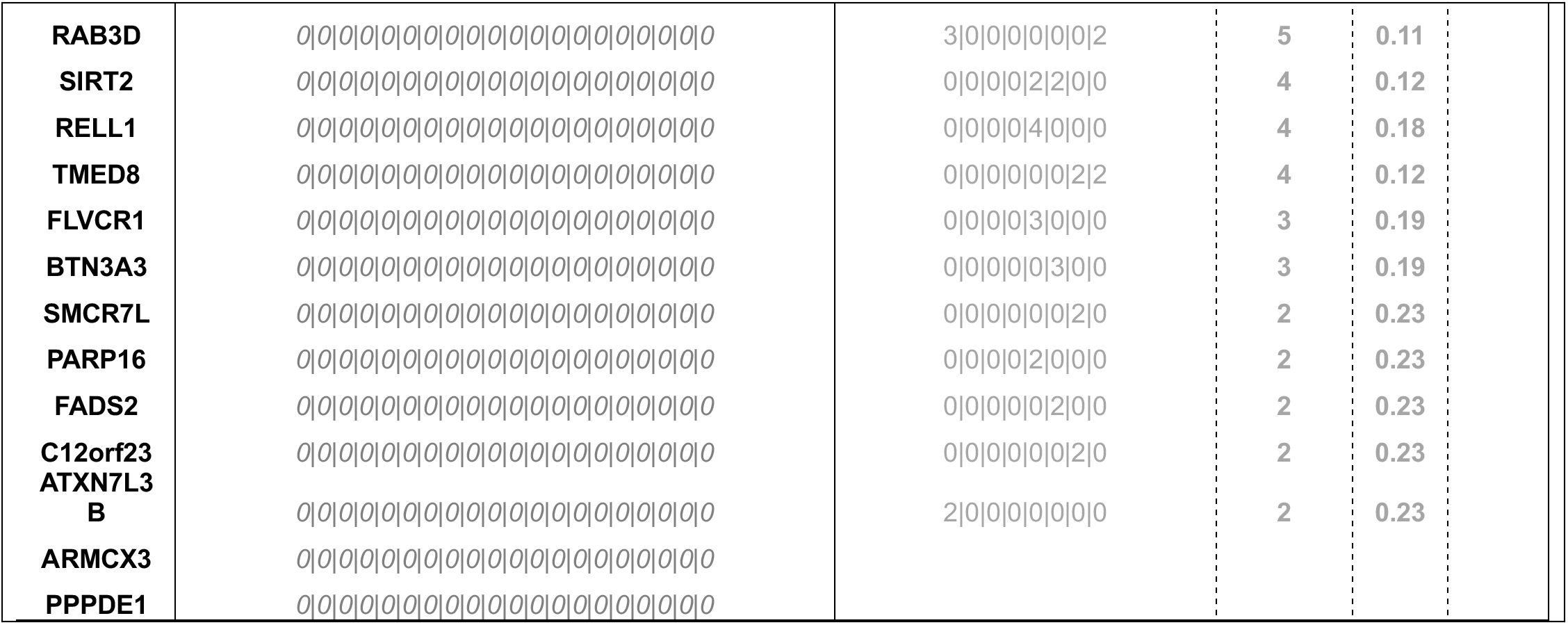
List of Bonafide BCL-2 Interactors in HEK293T Cells.

**Table 2.** Genotypic and Cytogenetic Characteristics of Patient Specimens.

| Patient Sample Number | Sex | Age | Diagnosis | Karyotyping | Mutations |
| --- | --- | --- | --- | --- | --- |
| AML 1 | F | 73 | De Novo AML | 45,XX,add(3)(q27),-7,add(8)(q24),t(9;22)(q34;q11.2)[4]/54,sl,+3,add(3)(q27),+7,+8,+10,+12,+13,+15,+20,+der(22)t(9;22)(q34;q11.2)[16] | N/A |
| AML 2 | M | 21 | De Novo AML | inv(16) (p13.1q22) | CKIT D816V |
| AML 3 | M | 79 | Prior MDS, AML Relapse | 46,XY,t(6;9) (p21;q34) | FLT3 ITD+ and FLT3 TKD mutation (D835); IDH2 R140Q; |
| AML 4 | M | 52 | De Novo AML | 45,XY,-7[3]/46,sl,+r(7)(p11q21)[11]/46,sdl1,der(5)t(1;5) (q31;p14)[5]/46,XY[1] | ASXL1 c.4120_4121insT; DNMT3A c.885delG; Notch1 c.4746_4747insTGGGGA; NRAS c. 182A>G |
| AML 5 | F | 49 | De Novo AML | Normal karyotype (46, XX) | FLT3 ITD+, FLT3 D835 |
| AML 6 | NA | NA | De Novo AML | NA | NA |
| AML 7 | M | 47 | Relapse AML | 46,XY,del(7)(q21)[8]/46,sl,del(5)(q31q35),add(12)(p13)[7]/46,sl,add(12)(p13),del(17)(q21)[3]/46,XY,del(9)(q22q32)[2] | IDH1 R132; CKIT D816V |
| AML 8 | F | 49 | Relapse AML | Normal karyotype (46, XX)[21] | FLT3 ITD+, FLT3 D835 |
| AML 9 | NA | 36 | Relapse AML | Normal karyotype | TET2.I1762V, ASXL1.L815P, CUX1.A448T, BCORL1.F111L, TP53.P33R |
| AML 10 | NA | NA | NA | clone 1: +8q, +3q, t(9;22); clone 2: clone 1 abnormalities AND +3, +10, +12 +13, +15 +20; clone 3: clone 2 abnormalities AND der (22); by FISH -7 | NA |
| AML 11 | F | 74 | De Novo AML | 46,XX[20] | FLT3 TKD (D835), NPM1+, TET2 c.4311_4318GAAAAAC, 41%, TET2.I1762V 24%, PTPN11.A72T 43%, ASXL1.L815P 100%, BCCORL1.F111L 100%, DNMT3AA.G354C 43%, CUX1.A448T 100%, NOTCH1.V2285I 41%, CBLC.P389S 38%. |
| AML 12 | F | 65 | Relapse AML | 46,XX,add(14)(q22)[4], 46,XX[16] | FLT3+, NPM1+, IDH1+ |
| AML 13 | F | 60 | De Novo AML | 46,XX,t(9;11)(p21;q23)[13]/47,sl,+21[4]/47,sl,+8[3] | KRAS.G13D 23%, KRAS.A146T 6%, PTPN11.T52A 47% |
| AML 14 | NA | NA | De Novo AML | 45,X,-Y; t(9;11)(p22;q23) | FLT3 ITD |
| AML 15 | NA | NA | NA | 46,XX,inv(3)(q21q26)[10] | NA |
| AML 16 | M | 52 | De Novo AML | 46,XY,add(1)(p11),del(5)(q15q33),del(7)(q22q36),der(11)t(1;11)(p31;p12-14)[20] , Loss of 5q31 and 7q31 | ASXL1 c.4120_4121insT; DNMT3A c.885delG; Notch1 c.4746_4747insTGGGGA; NRAS c. 182A>G |
| AML 17 | F | 77 | De Novo AML | 46,XX[20] | NPM1: W288fs 45%, RUNX1: S469P 6%, TET2: V1232del 48% |
| AML 18 | F | 80 | De Novo AML | 47,XX,+8[15]/47,XX,+11[1]/44-45,XX,add(5)(p12),dic(5;12)(q10;q10)t(11;14)(q13;q32),dic(17;18)(q10;q10),add(20)(q11.2)[cp8]/44,XX,dic(5;12)(q10;q10),der(8)(11qter-> | CBL M400R, SRSF2 P95H, TET2 Q916X and S1290X |
|  |  |  |  | 11q13::14q32- > 14q10::8p11.2- > 8qter),der(11)t(11;14)(q13;q32),dic(17;18)(q10;q10)[2]/46,XX[5] |  |
| AML 19 | M | 63 | De Novo AML | 46,XY[19] | FLT3 ITD+, NPM1+ |
| AML 20 | F | 64 | De Novo AML | NA | NA |
| AML 21 | M | 69 | CMML to AML | 46,XY[18] | NPM1.W288CFs*12 37%, ASXL1.G646Wfs*12 36%, NRAS.G12D 46%, STAG2.G46Rfs*41 11%, EZH2.Q533L 50%, EZH2.L149Q 49%, TET2.N1610Ifs*6 50%, TET2.Q1699* 50% |
| AML 22 | F | NA | De Novo AML | 46,XX,t(6;11)(q27;q23)[18]/92,slx2[2] | KRAS.G12C 16%, PTPN11.E76L 7% |
| AML 23 |  |  | Relapse AML | 46,XY[20] | NA |
| AML 24 | NA | NA | NA | Trisomy 8 in 37% of cells. Positive for IGH/CCND1 in 7.7% of cells, positive for trisomy 12 in 12.5% of cells, positive for trisomy 11 in 3.5% of cells, positive for a population with one copy of TP53 and three copies of centromere 17 comprising 3% of cells. | NA |

Based on the findings presented in this study, we propose the model shown in Fig. 6A, where sensitive LSCs require relatively low calcium due to inherently low basal respiration. Venetoclax treatment induces calcium overload through BCL-2 mediated reduction of SERCA, which disrupts OXPHOS and leads to cell death. Conversely, resistant LSCs (Fig. 6B) require relatively high calcium levels to support increased respiration. This state is maintained by decreased SERCA and increased MCU levels. Therefore, we postulate that venetoclax treatment has no effect on calcium flux in resistant specimens because SERCA is intrinsically suppressed and is therefore no longer a target for down-regulation in the context of BCL-2 inhibition. However, venetoclax-resistant LSCs are preferentially reliant on maintaining relatively high levels of mitochondrial calcium and are consequently sensitive to perturbations that inhibit calcium uptake (i.e., MCUi4, Ru265, and Mitox).

**Figure 6.**
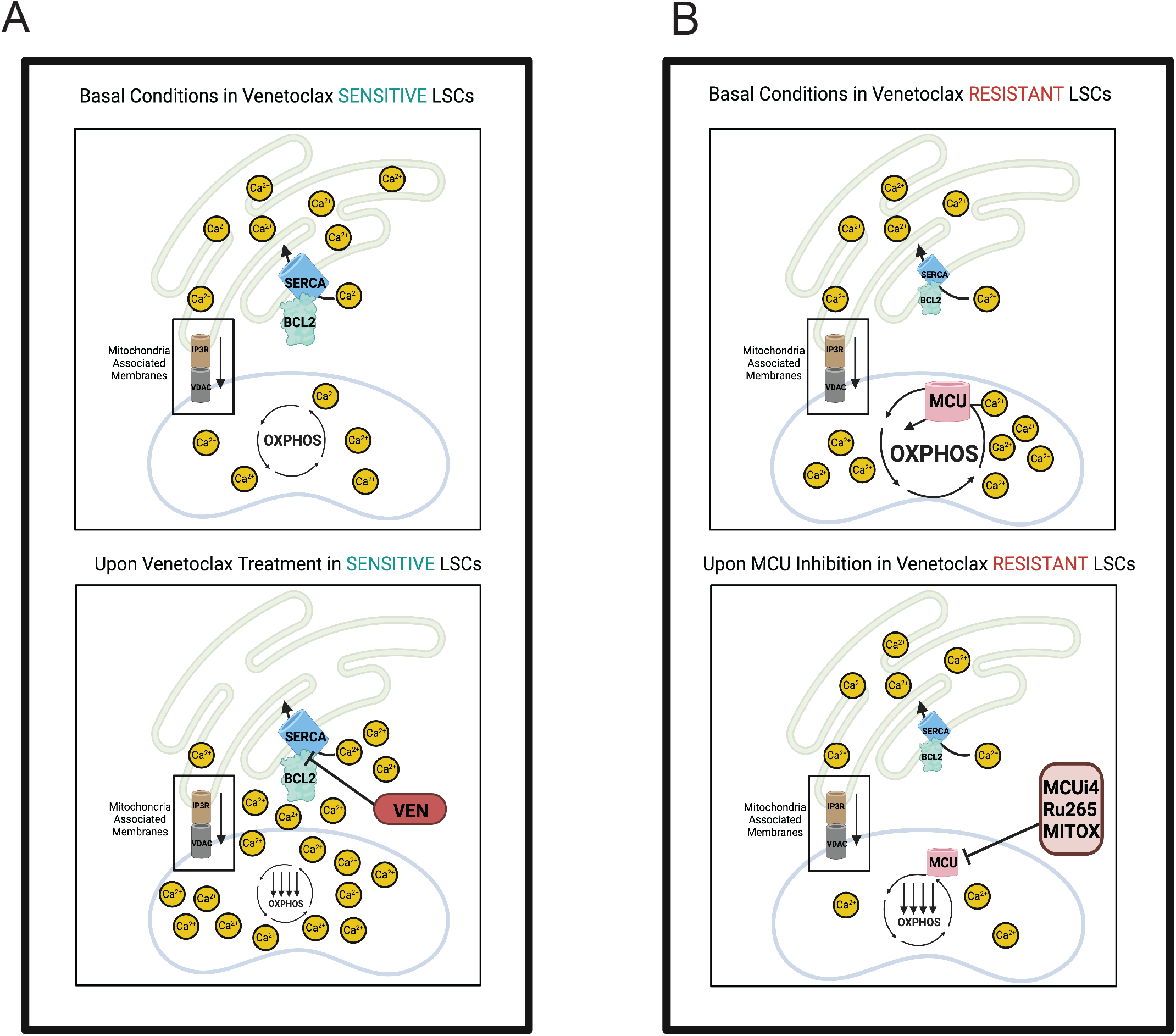
Hypothesized Model of Intracellular Calcium Dynamics in Venetoclax Sensitive versus Resistant LSCs. Hypothesized model of differences in intracellular calcium localization between venetoclax sensitive versus resistant LSCs. **A.** Due to inherently low basal respiration, venetoclax sensitive LSCs require relatively low basal mitochondrial calcium. During basal conditions, in venetoclax sensitive primary human AML LSCs, BCL2 promotes proper SERCA activity, thereby shuttling calcium into the ER for proper storage preventing cytosolic or mitochondrial calcium overload. MAMs facilitate proper exchange of calcium from the ER to mitochondria to fuel OXPHOS activity. Upon venetoclax treatment, SERCA activity it is decreased leading to decreased ER calcium uptake. Increased cytosolic calcium levels combined with transfer of calcium from MAMs leads to increased calcium being stored in the mitochondria leading to calcium overload and inhibition of proper mitochondrial metabolism. **B.** In contrast to sensitive LSCs, venetoclax resistant LSCs have evolved to have higher levels of mitochondrial calcium to support increased metabolic demands including higher OXPHOS activity. These functional changes are supported by decreased BCL-2 and SERCA3 levels concomitant with increased MCU expression. As a result, venetoclax treatment does not induce further decreased BCL-2 activity or SERCA levels. As venetoclax is unable to perturb intracellular calcium signaling in resistant LSCs, we hypothesized directly perturbing mitochondrial calcium content, a key ion required for OXPHOS activity, could target venetoclax resistant LSCs. Inhibition of MCU through genetic or pharmacologic inhibition leads to decreased mitochondrial calcium levels, OXPHOS activity and LSC activity. Taken together, our data demonstrate that either mitochondrial calcium overload or depletion is detrimental to mitochondrial metabolism and cell function.

## Discussion

We identify a novel function of venetoclax in modulating intracellular calcium localization that is directly linked to targeting of LSCs. Using transcriptomic analyses, we first established differences in expression of calcium mediated signaling genes between venetoclax-sensitive and resistant AML specimens. As LSCs are the key cell population responsible for disease initiation and progression, we focused subsequent functional studies on this population. Basal levels of mitochondrial calcium were significantly higher in resistant LSCs. These data support our previous findings which show distinct differences in the biology of venetoclax resistant LSCs that include increased basal mitochondrial metabolism and utilization of multiple fuel sources including fatty acids [5, 9]. In seeking to determine if and how venetoclax may impact this pathway, we analyzed the effects of treatment on sensitive versus resistant LSCs. We discovered that venetoclax treatment causes mitochondrial calcium overload subsequent to SERCA inhibition in sensitive LSCs only. This finding establishes for the first time the ability of venetoclax to perturb intracellular calcium dynamics and inhibit SERCA. In contrast, resistant LSCs did not undergo changes in mitochondrial calcium levels or SERCA activity upon venetoclax treatment. Therefore, we explored an alternative strategy to target venetoclax resistance which included inhibition of MCU, the only channel known to mediate uptake of calcium into the mitochondrial matrix. As calcium is required for the rate-limiting TCA cycle enzymes isocitrate dehydrogenase and alpha-ketoglutarate dehydrogenase, we hypothesized MCU inhibition would decrease OXPHOS activity, a key vulnerability of LSCs[5, 7–9] . Indeed, both pharmacologic and genetic inhibition of MCU led to decreased OXPHOS activity and LSC function in venetoclax resistant specimens. Lastly, we leveraged the recently described function of mitoxantrone as an MCU inhibitor to target venetoclax resistance. Importantly, at doses ∼10-100 fold lower than commonly used for chemotherapy, mitox efficiently inhibited OXPHOS and induced death of functionally defined LSCs. Taken together, these data demonstrate an unexpected and central link between BCL-2 and calcium signaling in the biological response to venetoclax in primitive AML cells. Additionally, our findings support clinical utility of mitox, specifically in the context of targeting venetoclax-resistant LSCs.

While previous reports have demonstrated that BCL-2 can have both positive and negative effects on SERCA structure and function [16, 43, 44], to our knowledge the data in Figure 2 are the first to indicate that BCL-2 may act to stabilize SERCA3 levels in primary human AML cells. Previous studies indicate that the BH4 domain is mainly responsible for the role of BCL-2 in regulating ER and mitochondrial calcium [17, 45–47]. As venetoclax was designed as a BH3 mimetic, it was not expected to influence calcium levels. Our data suggest a mechanism by which SERCA3 protein levels are rapidly reduced (∼3hrs) upon venetoclax binding of BCL2, leading to alteration of calcium homeostasis. Additionally, another mechanism of SERCA3 inhibition could be the oxidation of cysteine residues due to increased ROS levels post venetoclax treatment [2, 48]. It is also possible that the binding of venetoclax to BCL-2 causes a previously unrecognized steric hindrance of the BH4 domain leading to effects on SERCA3. A more detailed structural analysis of BCL2 interactions with SERCA3 appears warranted by our findings.

Aside from a role in regulating metabolism, several studies have documented the functional role of intracellular calcium homeostasis in AML cell lines and cancer stem cells [49–51]. For example, in AML cell lines mitochondrial calcium levels were increased in cells that evaded cytarabine treatment [49]. In addition, indirect evidence of a role for calcium signaling in LSCs was recently reported in studies that targeted ORP4L, a protein responsible for IP_3_ generation [51]. Specifically, pharmacological inhibition of ORP4L led to decreased cytosolic and mitochondrial calcium oscillations in LSCs which was associated with decreased OXPHOS activity and subsequent cytotoxicity [51]. These studies support our data showing intracellular calcium dynamics are crucial for LSC activity and can be leveraged for novel therapeutic strategies.

Our studies were motivated by the need to overcome venetoclax resistance in AML. This is a critical clinical need as venetoclax-based regimens have become ubiquitous in AML and most elderly patients are initially treated with these regimens. However, up-front resistance/relapse on therapy occurs in almost all patients. Therefore, we sought to investigate strategies to target venetoclax resistant LSCs that are clinically actionable. In particular, MCU inhibition was an attractive therapeutic strategy to target venetoclax resistance as previous studies have shown that MCU KO may not affect cells at basal conditions, but rather impair function during stressed situations [52]. Therefore, it is plausible that MCU inhibition is critical for malignancy but not necessarily for normal conditions. Our studies using normal hematopoietic stem/progenitor cells support this notion.

As there are currently no clinical grade MCU inhibitors, we explored previous agents that have been shown to inhibit mitochondrial calcium uptake. Of all FDA approved agents, mitox was the only inhibitor shown to lose its MCU inhibitory properties upon a single amino acid mutation strongly suggesting direct inhibition. Additionally, mitox is a well-established chemotherapeutic agent currently approved for the treatment of AML and other malignancies. Therefore, the safety and toxicity profile of this agent is well known. It is important to note that mitox is the only agent in this class (topoisomerase II inhibitors) that inhibits MCU due to its unique positively charged side chains [42]. Our data supported this as doxorubicin and etoposide, other topoisomerase II inhibitors, were unable to decrease mitochondrial calcium or colony formation in venetoclax-resistant specimens. Given these data, we tested mitox as a LSC targeting agent. Surprisingly, doses significantly lower than the plasma concentration of standard mitox regimens in AML patients (1uM), were effective at suppressing the colony formation of venetoclax resistant specimens [53]. Further, the doses used in our murine *in vivo* models are roughly equivalent to 1.5mg/m^2^/day based on morphometric calculations with previously established methods [54]. To our knowledge, these data are the first to show activity of mitox against LSCs. Based on our findings, we postulate that lower doses of mitox may be effective in venetoclax resistant AML which may be particularly useful in the context of treating elderly patients with comorbidities. Our data suggest that lower dose mitox alone may not be sufficient to reduce bulk tumor disease in florid venetoclax relapse scenarios. Therefore, we are currently designing clinical trials to use lower dose mitox as an agent to target MRD+ disease, thereby delaying time to relapse or progression of disease. Indeed, mitox is a well-established agent in the AML field and has historically had limited efficacy in the chemotherapy relapsed/refractory setting. Currently, mitox is being used in conjunction with other agents in clinical trials for both upfront and chemotherapy relapse/refractory AML (NCT03839446, NCT05522192, NCT04195945, NCT04797767, NCT04330820, NCT03531918). However, the effects of mitox on venetoclax-resistant disease have never been clinically tested. Additionally, considerable evidence suggests that venetoclax-resistant LSCs have unique features that make them biologically distinct from chemotherapy-resistant LSCs [5, 9, 55]. Our data show that hyper-sensitivity to mitox is one such feature, thereby indicating that clinical evaluation is warranted specifically in the context of venetoclax-resistant AML.

Beyond AML, our findings are relevant to other malignancies. Indeed, perturbation of intracellular calcium channels in the ER and mitochondria such as IP3R, MCU, and SERCA led to decreased cancer stem cell activity in breast cancer, glioblastoma, and melanoma stem cells, suggesting a central role for calcium in cancer stem cell function [50]. Given the rapid decrease in SERCA protein levels observed following venetoclax treatment, we suggest that other tumor types with aberrant SERCA expression may also be responsive to venetoclax (e.g., prostate, breast, colon and T-cell acute lymphoblastic leukemia) via a similar mechanism [56]. For instance, in T-cell acute lymphoblastic leukemia, SERCA inhibition has been shown to target NOTCH1 mutated disease [57]. However, there are currently no clinically available SERCA or NOTCH1 inhibitors. Therefore, venetoclax may be an alternative strategy to target SERCA in this context.

In conclusion, we demonstrate that venetoclax responsiveness is associated with unique calcium biology properties. Surprisingly, venetoclax was able to induce significant changes in mitochondrial calcium levels and inhibit SERCA3 levels in sensitive LSCs only. Further, venetoclax-resistant LSCs have adapted to their unique metabolic demands by increasing basal mitochondrial calcium levels and do not undergo changes in calcium signaling upon treatment. Genetic and pharmacologic inhibition of MCU led to decreased OXPHOS activity and LSC targeting. These data establish calcium biology as a critical mediator of LSC survival as either mitochondrial calcium overload or insufficiency targets venetoclax sensitive or resistant LSCs, respectively. Lastly, we postulate mitoxantrone may be an effective component of anti-LSC therapy specifically in the context of venetoclax-resistant AML.

## Methods

### Human Specimens

Primary human AML specimens were obtained by apheresis product, peripheral blood or bone marrow. Mobilized peripheral blood was obtained from normal healthy donors. All patients gave informed consent for procurement of samples on the University of Colorado tissue procurement protocol. The University of Colorado Institutional Review Board approved the retrospective analysis. Further details on each human AML specimen used for analysis are included in Supplementary Table 1.

### Human Specimen Culturing

Primary human AML specimens were cultured as previously described using InVitria Metabolomics AF media supplemented 10nM human cytokines SCF (PEPROTech, 300-07), IL3 (PEPROTech, 200-03), and FLT3 (PEPROTech, 300-19) [9]. In addition, the media was supplemented with low density lipoprotein (Millipore, 437744) and penicillin/streptomycin.

### Cell Sorting

Primary human AML specimens were sorted for ROS-low LSCs as previously described [8, 34]. Briefly, specimens were thawed and stained with DAPI (EMD Millipore, no. 278298; dilution 500nM) to exclude dead cells, CD19 (BD, no. 555413; dilution 1:20) and CD3 (BD, no. 557749; dilution 1:40) to exclude lymphocytes, CD45 (BD, no. 571875; dilution 1:40) to identify the blast population and CellROX deep red (Thermo Fisher, no. C10422; dilution 5uM) to identify the 20% of AML blasts with the lowest ROS stain signal, deemed “ROS-Low LSCs”.

### Mitochondrial Calcium and Membrane Potential Measurement by Flow Cytometry

100,000-500,000 primary human AML Ros-Low cells or MOLM-13 cells were used for analysis of mitochondrial calcium content or mitochondrial membrane potential. For mitochondrial calcium measurements, cells were incubated with Rhod2AM (Invitrogen, R1245MP; 500nM) for 30 minutes at 37°C. Cells were then washed with calcium free, magnesium free 1X PBS (Corning, 21-031-CV) twice and resuspended in 1X calcium free, magnesium free PBS supplemented with 2% FBS (Atlas Biologicals, no. F-500-D) and analyzed by flow cytometry (PE channel). For co-localization studies using confocal microscopy of Rhod2AM with mitochondria, 20,000 cells were used per condition. Cells were fixed with 4% PFA for 20 minutes at 4 degrees and permeabilized with 0.1% Triton X100 for 15 minutes at room temperature. Slides were mounted using Prolong Gold Antifade mounting media (Thermo Fisher Scientific, P36935) containing DAPI. The stained cells were analyzed by a LSM 780 confocal microscope system (Carl Zeiss) equipped with an inverted microscope using a Plan Apochromat x40 H2O immersion lens. Images were analyzed and processed using Image J and Adobe Photoshop v23.4.1. Detection for Rhod2AM was done on 552/581 channel, MitoTracker was done on GFP channel and detection of nucleus was done using DAPI. Rhod2AM was labeled as described above and MitoTracker was stained as we previously established in ROS-low cells [58]. For mitochondrial membrane potential measurements, cells were incubated with TMRE (Invitrogen, T669; 1uM) for 30 minutes at 37°C. Cells were then washed with calcium free, magnesium free 1X PBS twice and resuspended in 1X calcium free, magnesium free PBS supplemented with 2% FBS and analyzed by flow cytometry (PE channel).

### Viability Measured by Flow Cytometry

Cell viability was measured using Dapi and measured by flow cytometry. Cells were stained in PBS supplemented with 2% FBS.

### Seahorse

The extracellular flux assay kit XF96 (Agilent Technologies, no.102417-100) was used to measure OCR per manufacturer’s instructions and as previously described [5]. Briefly, 200,000-500,000 cells were used per well per condition with 5 technical replicates per condition and measured using the MitoStress Test as per manufacturers protocol. Cells were plated in Cell-Tak-coated (Corning, no. 354240) XF96 cell culture microplates. OCR was measured at basal levels and after injections of oligomycin A (Sigma, no. 75351; 5ug/mL), FCCP (Sigma, no. C2920; 2umol/L) and Antimycin A with Rotenone (Sigma, no. A8674 and no. R8875, 5umol/L each).

### Enzyme Activity assays

Activity of either isocitrate dehydrogenase or alpha-ketoglutarate dehydrogenase was measured by manufacturer’s protocol (Abcam, no. ab102528 or no. ab185440). 500k cells were used per replicate per condition and assay was run in technical triplicate per condition.

### BioID Sample Preparation

BCL-2 open reading frame was cloned in-frame with an N-terminus Flag-BirA R118G (Flag-BirA*) into a tetracycline-inducible pcDNA5 FLP recombinase target/tetracycline operator (FRT/TO) expression vector. Flp-In T-REx HEK293 cells were transfected with the FlagBirA*-BCL-2 or control FlagBirA* only. Cells were incubated with 1µg/mL tetracycline (Sigma-Aldrich) and 50 µM biotin (BioShop) in DMEM supplemented with 10% FBS and 1% pen/strep for 24 hrs at 37°C with 5% CO_2_. Cells were collected, rinsed in PBS and lysed in RIPA buffer. The lysates were sonicated twice for 10 sec at 35% amplitude (Sonic Dismembrator 500; Fischer Scientific) and centrifuged at 16,000 rpm for 30 min at 4°C. Supernatants were then passed through Micro Bio-Spin Chromatography column (Bio-Rad) and incubated with high-performance streptavidin sepharose (GE Healthcare) for 3 hours at 4°C on an end-over-end rotator. Beads were washed 6 times with 50 mM ammonium bicarbonate and then treated with TPCK-treated modified trypsin (Promega) for 16 hours at 37°C on an end-over-end rotator. Supernatants were lyophilized and desalted using C18 tips prior to downstream MS analysis.

### Liquid chromatography – Mass Spectrometry (LC-MS) for Bio-ID

Lyophilized samples reconstituted in 0.1% HCOOH were loaded on a pre-column (C18 Acclaim PepMap^TM^ 100, 75µM x 2cm, 3µm, 100Å) prior to chromatographic separation through an analytical column (C18 Acclaim PepMap^TM^ RSLC, 75µm x 50cm, 2µm, 100Å) by HPLC over a reversed-phase gradient (120-minute gradient, 5-30% CH_3_CN in 0.1% HCOOH) at 225nL/min on an EASY-nLC1200 pump in-line with a Q-Exactive HF (Thermo Scientific) mass spectrometer operated in positive ESI mode. An MS1 ion scan was performed at 60,000 fwhm followed by MS/MS scans (HCD, 15,000 fwhm) of the 20 most intense parent ions (minimum ion count of 1000 for activation). Dynamic exclusion (10 ppm) was set at 5 seconds. To identify peptides and proteins, raw files (.raw) were converted to .mzML format using Proteowizard (v3.0.19311). Peak list files were searched using X!Tandem (v2013.06.15.1) and Comet (v2014.02.rev.2) against the human RefSeqV104 database (36,113 entries). Search parameters specified a parent ion mass tolerance of 15 ppm and an MS/MS fragment ion tolerance of 0.4 Da, with up to two missed cleavages allowed for trypsin. No fixed modification was set. Deamidation (NQ), oxidation (M), acetylation (protein N-term) were set as variable modifications. Data were processed through the trans-proteomic pipeline (TPP v4.7) using iProphet. Proteins were identified with an iProphet cut-off of 0.9 and at least two unique peptides. High-confidence proximity interactors were identified using Significance Analysis of INTeractome (SAINT,) [59] comparing FlagBirA*-only samples to FlagBirA*-BCL-2 samples using a Bayesian false discovery rate (BFDR) cut-off of ≤0.01 (1%). All mass spectrometry data have been deposited in the MassIVE repository (massive.ucsd.edu) under accession MSV000090632.

### Immunoblotting

Protein lysates were loaded on a polyacrylamide gel and transferred to a polyvinylidene difluoride membrane using the mini trans-blot transfer system (Biorad). To probe for specific antigens, blots were probed with primary antibodies of the following targets: BCL-2 (Cell Signaling Technologies, no.15071), Actin (Cell Signaling Technologies, no.4970), SERCA3 (Biorad, no. VPA00530) and MCU (Cell Signaling Technologies, no.14997) overnight at 4°C on a shaker. All primary antibodies were used at a 1:1000 dilution. After overnight incubation, membranes were incubated with respective HRP conjugated secondary antibodies (Biorad) for 1 hour at room temperature and imaged by the ChemiDoc Imaging System (Biorad). Images were analyzed and processed using Image Lab and quantitation of band intensity was done through Image J.

### siRNA Transfection of Primary Human AML Specimens

Primary human AML specimens were transfected with siRNA using previously established protocols [5, 60]. Briefly, 2×10^5^ cells were electroporated in Buffer T containing 50nM siRNA, final concentration, targeting either a scrambled nontargeting sequence, BCL-2, ATP2A3 or MCU (Dharmacon, no.D-001810-01, no.L-003307-00, no.L-006114-00, no.L-015519-02).

### Proximity Ligation Assays

Proximity ligation assays (PLA) were performed according to manufacturer’s protocol for both flow cytometry and in situ confocal microscopy detection (Sigma, DUO94104 and DUO92013 respectively). For flow cytometry, 100,000 cells were used per reaction. Fixation was done with 4% PFA for 15 minutes at room temperature and permeabilization was done for 10 minutes with ice cold methanol. For in situ detection through confocal microscopy, 20,000 cells were used per condition. Cells were fixed with 4% PFA for 20 minutes at 4 degrees and permeabilized with 0.1% Triton X100 for 15 minutes at room temperature. Slides were mounted using Prolong Gold Antifade mounting media (Thermo Fisher Scientific, P36935) containing DAPI. The stained cells were analysed by a LSM 780 confocal microscope system (Carl Zeiss) equipped with an inverted microscope using a Plan Apochromat x40 H2O immersion lens. Images were analyzed and processed using Image J and Adobe Photoshop v23.4.1. Detection for PLA signal was done on FarRed channel and detection of nucleus was done using DAPI. Primary antibodies used include BCL-2 (Cell Signaling Technologies, no.15071), SERCA3 (Biorad, no. VPA00530), IgG Mouse (Cell Signaling Technologies, no. 5415S), IgG Rabbit (Cell Signaling Technologies, no. 3900S), VDAC1 (Abcam, no. 15895) and IP3R1 (Santa Cruz, no. 271197). Antibody incubation was done overnight at 4° C with dilution factor 1:100. Amplification step was done overnight at 37 ° C.

### Gamma H2AX Assays

Briefly, gamma-h2AX levels were measured for conditions indicated in figures using manufacturer’s protocol (Sigma, no. 17-344). 200k cells were used per replicate per condition and assay was run in technical triplicate per condition.

### Colony Forming Assays

Primary AML specimens or normal mobilized peripheral blood samples were plated in human methylcellulose (R&D systems) at cell concentrations indicated in respective figures. Samples were treated with small molecule inhibitors for 16 hours, washed of drug and added into methylcellulose cultures. Colonies were counted at 2 weeks after initial plating.

### Engraftment Assays

Leukemia stem cell function was assessed by measuring engraftment of primary AML specimens or mobilized peripheral blood samples from health donors with indicated therapies transplanted into NSGS mice. One day prior to transplant, freshly thawed primary AML cells were treated in culture dishes overnight with indicated agents in media with cytokines as listed above. NSG-S mice were conditioned with 25 mg/kg busulfan via i.p. injection. Second day at injection, overnight-treated primary AML cells were washed and resuspended in calcium free, magnesium free PBS supplemented with FBS. Anti-human CD3 antibody (BioXCell) was added at a final concentration of 1 μg/10^6^ cells to avoid potential graft-versus-host disease. Per mouse, 2 × 10^6^ cells in 0.1 mL saline were tail vein injected; there were 8 to 16 mice per experiment group. Mice engrafted with primary AML cells were sacrificed after 4 to 8 weeks. Engraftment was measured by flow cytometry for human CD45+ cells (BD no. 571875, dilution 1:100). All animal studies were done at the University of Colorado under Institutional Animal Care and Use Committee– approved protocol no. 308. The University of Colorado is accredited by the Association for Assessment and Accreditation of Laboratory Animal Care (ALAC), abides by the Public Health Service (PHS) Animal Assurance of Compliance and is licensed by the United States Department of Agriculture.

### In Vivo PDX Drug Treatments

Effects of mitoxantrone on PDX tumor burden were assessed as follows. One day prior to transplant, NSG-S mice were conditioned with 25 mg/kg busulfan via i.p. injection. Second day at injection, primary AML cells were washed and resuspended in calcium free, magnesium free PBS supplemented with FBS. Anti-human CD3 antibody (BioXCell) was added at a final concentration of 1 μg/10^6^ cells to avoid potential graft-versus-host disease. Per mouse, 1 × 10^6^ cells in 0.1 mL saline were tail vein injected; mouse number per experiment per condition indicated in respective figure legends. Mice were treated with indicated condition (PBS-vehicle or mitoxantrone) when at least 20% bone marrow disease burden was present. Treatment regimen was once per day, 4 days in a row of either PBS or mitoxantrone (0.5mg/kg/day). For secondary transplants, bone marrow cells harvested from the previous experiment were injected in equal numbers into NSG-S mice conditioned with busulfan as described above (1 million cells/mouse per condition). Mice were sacrificed after 4 weeks. Engraftment was measured by flow cytometry for human CD45+ cells (BD no. 571875, dilution 1:100). All animal studies were done at the University of Colorado under Institutional Animal Care and Use Committee– approved protocol no. 308. The University of Colorado is accredited by the Association for Assessment and Accreditation of Laboratory Animal Care (ALAC), abides by the Public Health Service (PHS) Animal Assurance of Compliance, and is licensed by the United States Department of Agriculture.

### Quantitative RT-PCR

RNA was isolated using the RNeasy plus mini kit (QIAGEN) following manufactures instructions. cDNA was synthesized using qScript cDNA SuperMix as per manufacturers protocol (Quanta Bio, no. 95048). Quantitative real-time PCR was performed with LightCycler96 real-time PCR using PerfeCTA SYBR Green FastMix (Quanta Bio, no. 95072-05K). Following primers were used: ACTIN (For: GGACTTCGAGCAAGAGATGG Rev: AGCACTGTGTTGGCGTACAG), GAPDH (For: TGTGGGCATCAATGGATTTGG Rev: ACACCATGTATTCCGGGTCAAT), BCL-2 (For: GGTGGGGTCATGTGTGTGG Rev: CGGTTCAGGTACTCAGTCATCC), SERCA3 (For: GGAACCACATGCACGAAGAA Rev: TGAGGTACACACCGGAGACT), and MCU (For: AGAGACTGAGAGACCCATTACA Rev: GTTCCTTCTGCCAGGATTCA).

### DE Analysis of Bulk RNA-seq Dataset

Raw fastqs were obtained from GEO (GSE132511) [5]and processed using salmon v0.10.2. Data was imported into R using tximport. For principal component analysis, the data was batch corrected using limma v3.46 and variance stabilized using DESeq2 v1.30.1. Differential expression analysis was performed using DESeq2 comparing sensitive to resistant and correcting for batch effect using the Wald test. Significantly differentially expressed genes were defined as absolute log2 fold change >= 0.5 & adjusted pvalue < 0.05. Log2 fold change values reflect sensitive/resistant. There were 1953 DEGs with negative log2 fold change values (up in resistant) and 1984 DEGs with positive log2 fold change values (up in sensitive).

### GSEA Analysis

Gene set enrichment analysis was performed using the fgsea v1.16 package and GO biological process pathways were downloaded from BROAD (https://www.gsea-msigdb.org/gsea/msigdb/). Ranked test statistics from the above described DE test was used as the input to fgsea. Ten thousand permutations were used to calculate significance and enrichment scores. Significant pathways were those with adjusted p values < 0.05.

### CITE-SEQ Analysis

Raw sequencing data for gene expression, antibody derived tag (ADT; surface protein), and hashing libraries were processed using STARsolo 2.7.8a (https://doi.org/10.1093/bioinformatics/bts635; https://github.com/alexdobin/STAR/blob/master/docs/STARsolo.md) with the 10X Genomics GRCh38/GENCODE v32 genome and transcriptome reference (version GRCh38_2020A; https://support.10xgenomics.com/single-cell-gene-expression/software/release-notes/build#GRCh38_2020A) or a TotalSeq barcode reference, as appropriate. Hashed samples were demultiplexed using GMM-Demux (https://doi.org/10.1186/s13059-020-02084-2; https://github.com/CHPGenetics/GMM-Demux). Next, cell-containing droplets were identified using dropkick 1.2.6 (https://doi.org/10.1101/gr.271908.120; https://github.com/KenLauLab/dropkick) using manual thresholds when automatic thresholding failed, ambient RNA was removed using DecontX 1.12 (https://doi.org/10.1186/s13059-020-1950-6; https://github.com/campbio/celda) and cells estimated to contain >50% ambient RNA were removed, and doublets were identified using DoubletFinder 2.0.3 (https://doi.org/10.1016/j.cels.2019.03.003; https://github.com/chris-mcginnis-ucsf/DoubletFinder) and removed. Remaining cells were then filtered to retain only those with > 200 genes, 500-80,000 UMIs, < 10-20% of UMIs from genes encoded by the mitochondrial genome (sample dependent based on UMI distributions), < 5% of UMIs derived from HBB, < 20,000 UMIs from antibody-derived tags (ADTs), and >100-2,750 UMIs from antibody derived tags (sample dependent based on UMI distribution). Filtered cells were modeled in latent space using TotalVI 0.18.0 (ttps://doi.org/10.1038/s41592-020-01050-x; https://github.com/scverse/scvi-tools) to create a joint embedding derived from both RNA and ADT expression data, corrected for batch effects, mitochondrial proportion, and cell cycle. Scanpy 1.8.2 (https://doi.org/10.1186/s13059-017-1382-0; https://github.com/scverse/scanpy) was used to cluster the data in latent space using the leiden algorithm (https://doi.org/10.48550/arXiv.1810.08473) and marker genes were identified in latent space using TotalVI. Clusters were annotated using clustifyr 1.9.1 (https://doi.org/10.12688/f1000research.22969.2; https://github.com/rnabioco/clustifyr) and the leukemic/normal bone marrow reference dataset presented in (https://doi.org/10.1038/s41590-021-01059-0). Scanpy and Seurat 4.1.1 (https://doi.org/10.1016/j.cell.2021.04.048; https://github.com/satijalab/seurat) were then used to generate UMAP projections from the TotalVI embeddings and perform exploratory analysis, data visualization, etc. Antibodies used are in Table 3.

**Table 3.** CiteSeq Antibodies Used for Clustering.

CITESEQ- Total Seq B Cat Antigen Clone
|  |
| --- |
| 300565 CD4 RPA-T4 |
| 343129 CD7 CD7-6B7 |
| 362561 CD56 5.1H11 |
| 344853 CD3 SK7 |
| 302263 CD19 HIB19 |
| 366635 CD33 P67.6 |
| CD34 581 |
| 328147 CD90 5E10 |
| 313247 CD117 104D2 |
| 304161 CD45RA HI100 |
| 306047 CD123 6H6 |
| 301857 CD14 M5E2 |
| 307661 HLA-DR L243 |
| 301357 CD11b ICRF44 |
| 305049 CD64 10.1 |
| 303547 CD38 HIT2 |
| 368547 CD45 2D1 |
| 304066 CD45 HI30 |
| 336233 CD36 5-271 |
| 336125 CD93 VIMD2 |
| 312123 CD9 HI9a |
| 355125 CD70 113-16 |
| 302851 CD27 O323 |

## Statistical Analysis

Statistical analyses of all biological assays were performed utilizing denoted tests in Graphpad Prism v.9.0. Graphs and data are visualized as noted in figure legends. In all figures, ns indicates a not significant *P* value of >0.05; *, **, ***, and **** indicate *P* < 0.05, *P* < 0.01, *P* < 0.001, and *P* < 0.0001, respectively. Animal experiments were carried out based on previous power analysis and publications regarding AML xenograft models from groups utilizing the Lenth power calculator, and animal sample sizes of eight or greater based on expected standard deviation for a two-sample t-test. Technical replicates from Seahorse experiments were excluded when outliers (pmolmin^-1^ readings for OCR) were identified through an outlier analysis using Grubb’s test or when negative values occurred as indicated by manufacturers protocol for analysis. Seahorse experiments were done in technical replicates of five and multiple specimens from patients to account for technical issues with plates and collection of values. Randomization was applied in animal experiments as animals were randomized to injection and treatment groups. Investigators were not blinded to allocation during experiments and outcome assessment.

## Data Availability Statement

All sequencing data have been deposited into public database. The CITE-Seq data can be found at the GEO database and are available via accession number GSEXXXXXX.

## Acknowledgements

AIS is supported by the NCI NRSA F30 grant NCI (F30CA254251). MJA is supported by the NCI F32 award (F32CA275350). MLA is supported by the BL&CS Career Development Award (1IK2BX005603-01A1). AW is supported by American Cancer Society (CSDG-22-018-01-CDP), Morgan Adams Foundation (R6251I), and Swim Across America. CJ is supported by the LLS Special Fellow program. DAP is supported by the Leukemia and Lymphoma Society’s Scholar in Clinical Research Program, the Robert H. Allen MD Chair in Hematology Research and the V Foundation Clinical Scholar Program. BMS is supported by the V Foundation Clinical Scholar Program. CTJ is generously supported by the Nancy Carroll Allen Chair in Hematology Research, a Leukemia and Lymphoma Society SCOR grant (7020-19), NIH R35CA242376, and Veterans Administration merit award BX004768-01. We would also like to acknowledge the Colorado Nutrition Obesity Research Center (NIH P30 DK48520) for providing core for seahorse assays.

## Authorship Contributions

AIS and CTJ conceived the project and wrote the manuscript. AIS, HT carried out the studies and data analysis. KE, SSP, BMS, AG, MR and SS performed CITE-Seq experiments and subsequent analysis. JA performed confocal microscopy. AK and TY assisted with all PDX experiments. SSP, MLA, SBP, MM, AW, IS and RM assisted with molecular, genetic and pharmacological studies. JSG, CL, CJ and BR performed and analyzed BIO-ID experiments. CS, DAP and BMS assisted with clinical analysis and patient sample acquisition.

---

**Supplementary Figure 1.**
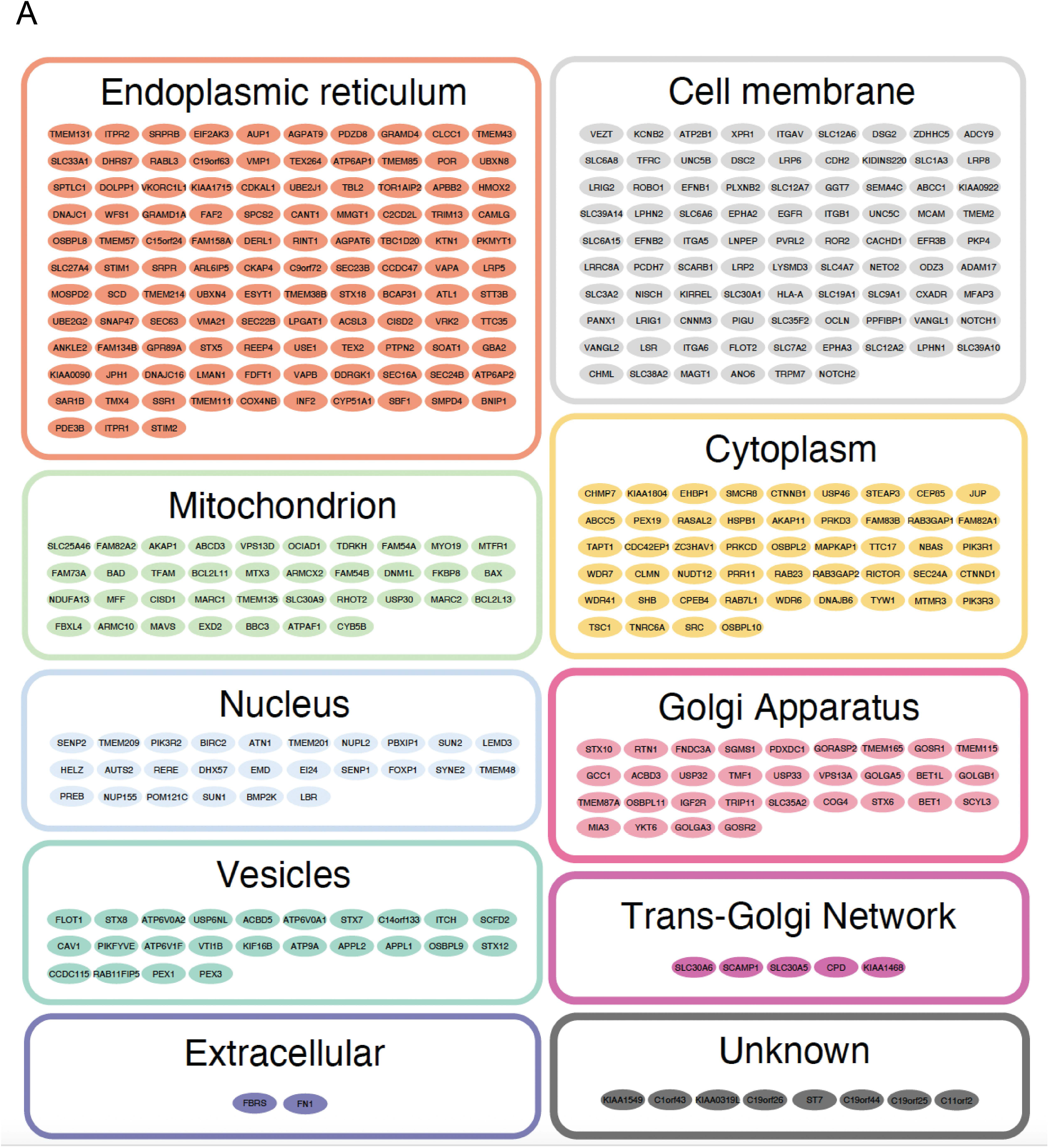
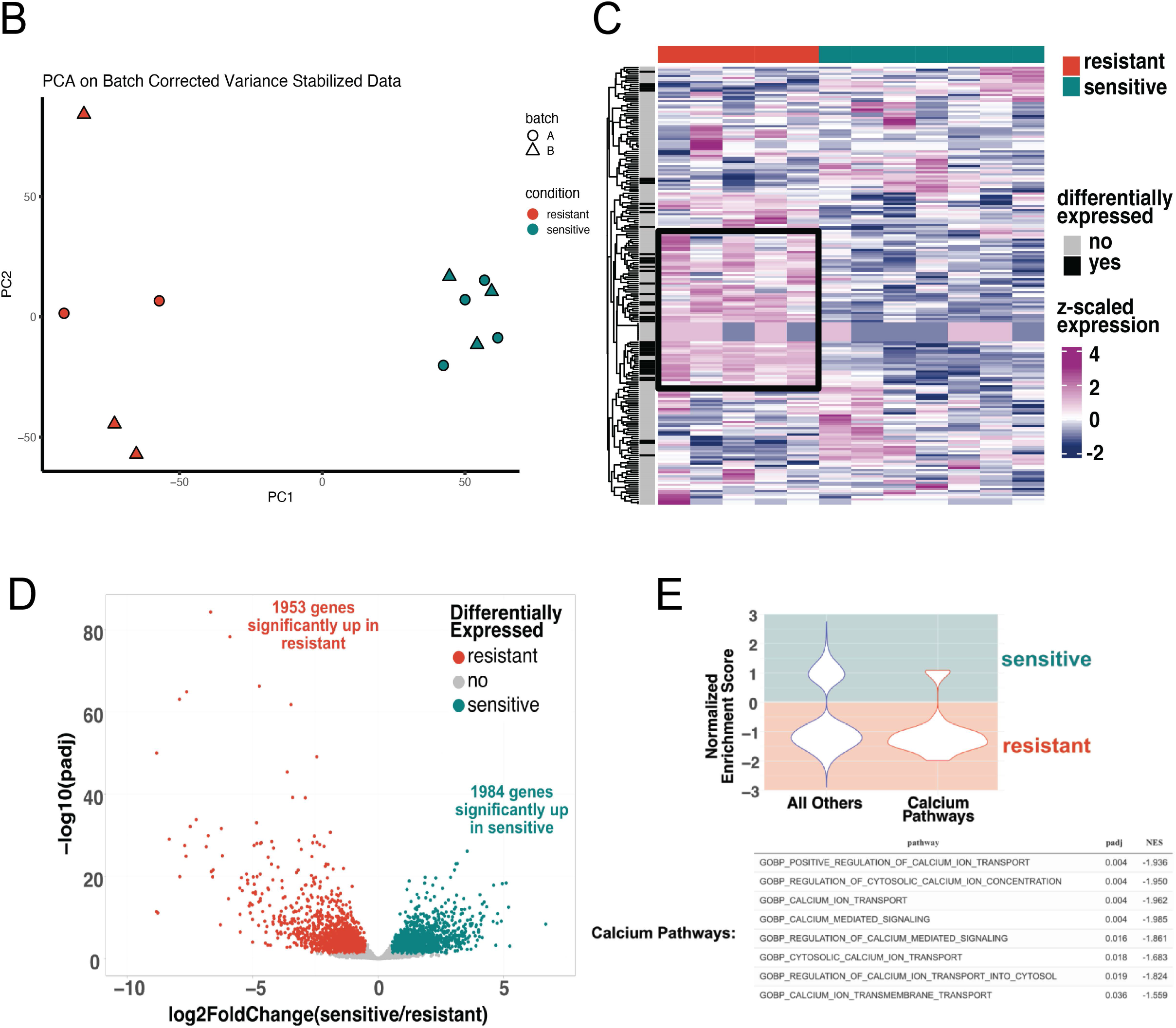
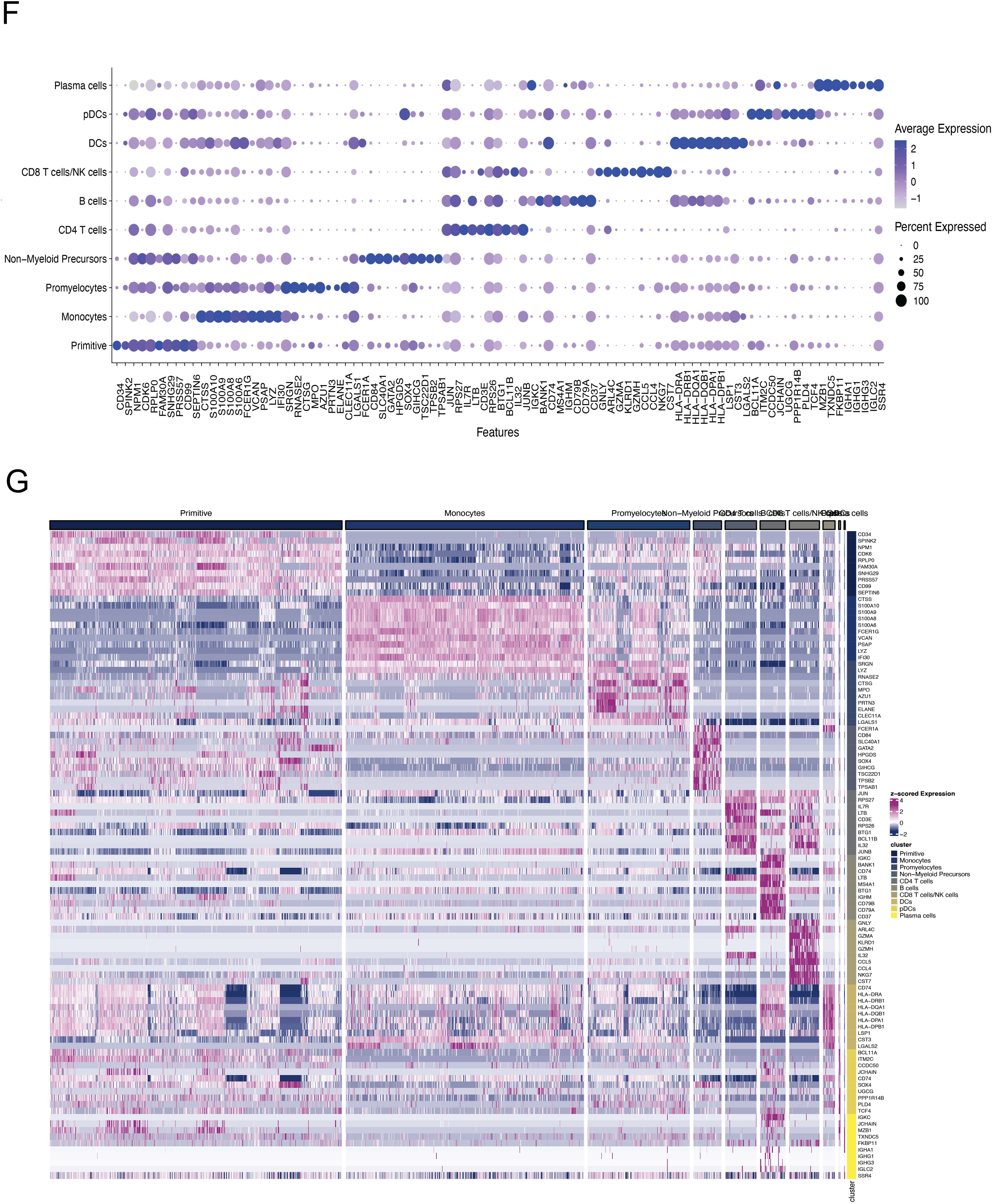
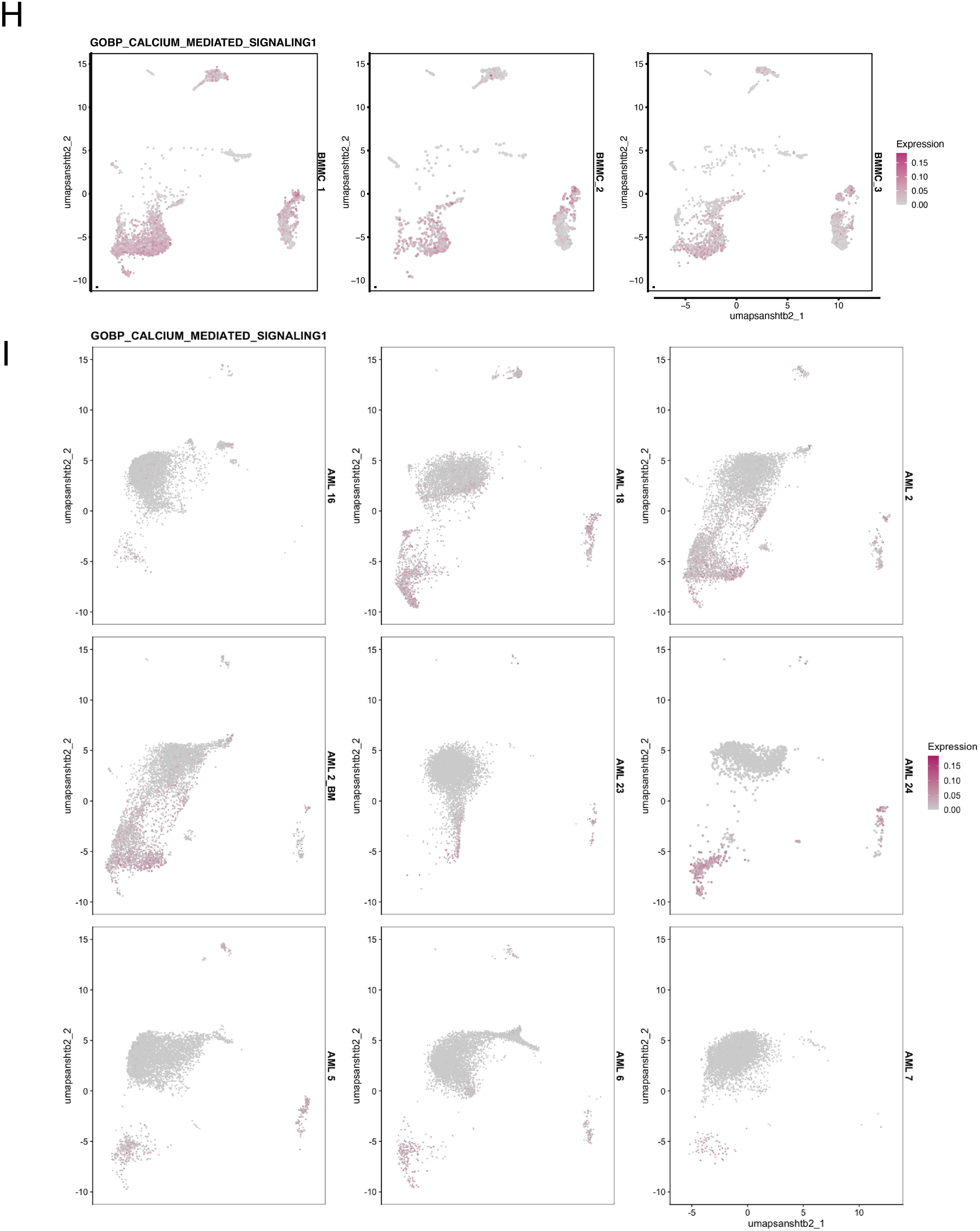
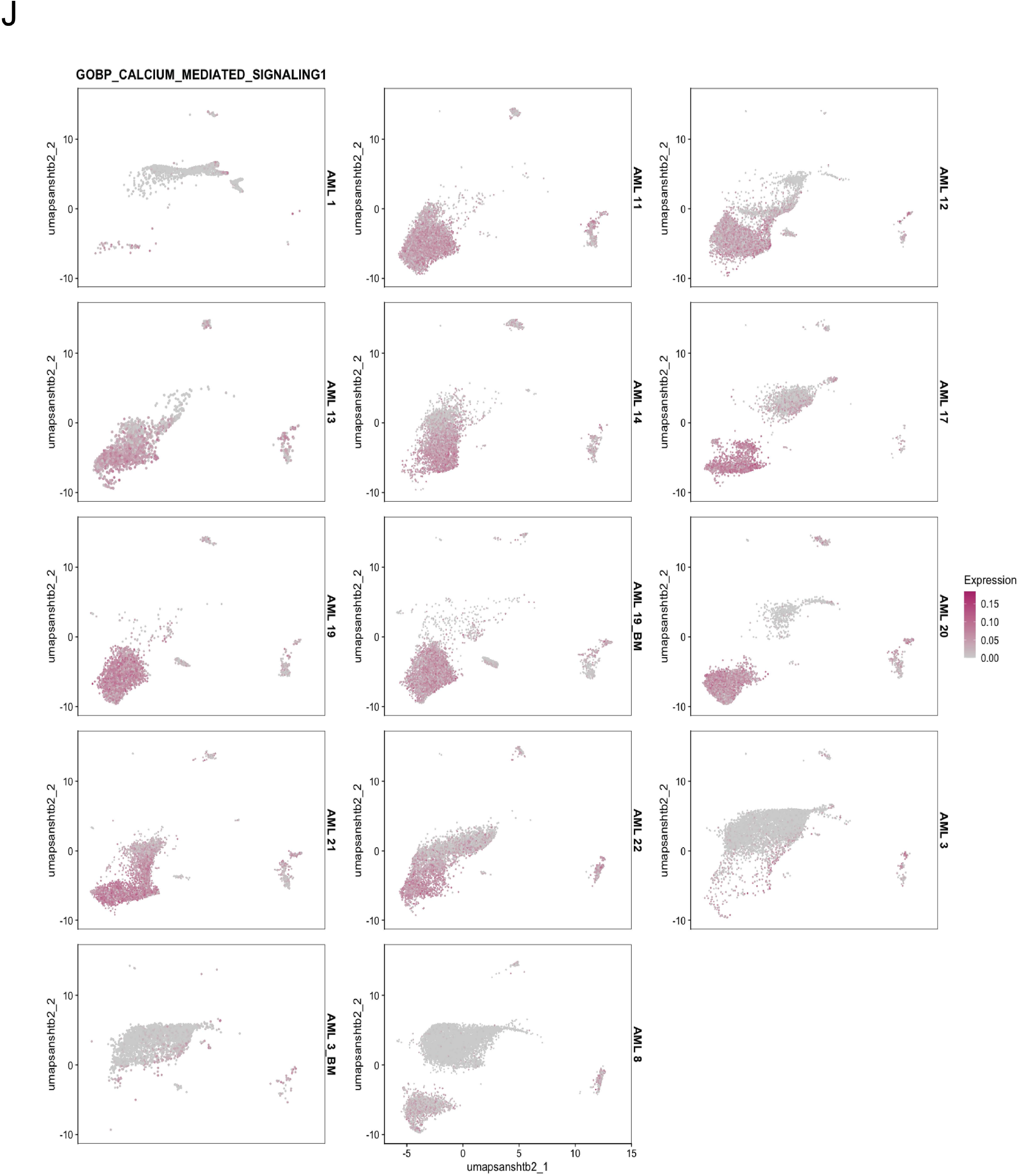
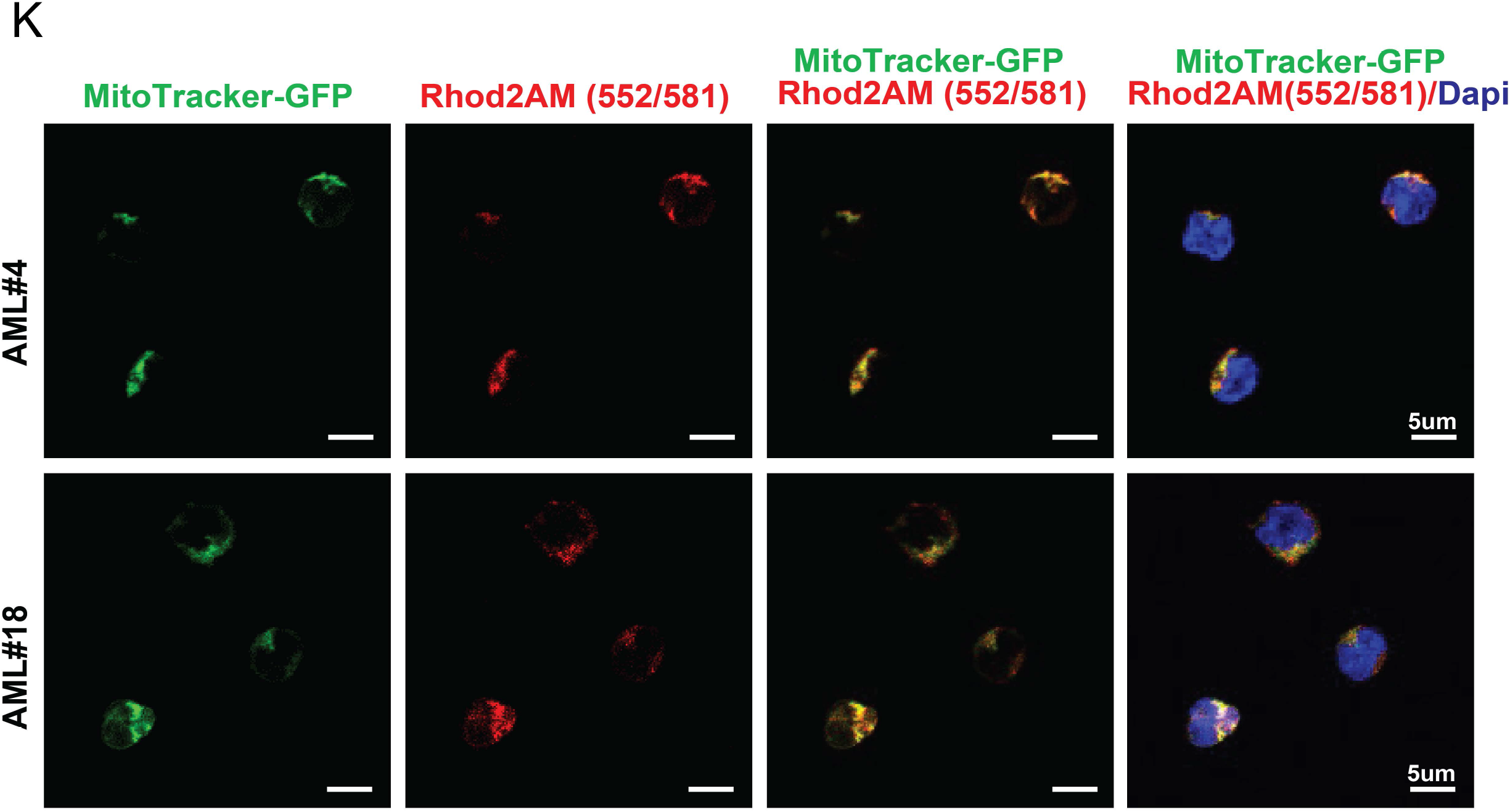
Venetoclax Responsiveness is Associated with Intrinsic Differences in Calcium Pathway Signaling. **A.** BCL-2 interacts with signaling proteins using BioID in T-REx HEK293 cells. All BCL-2 proximity interactors organized by established cellular location. **B-E.** Bulk RNA-sequencing performed in primary human AML specimens sorted for ROS-low cells that were either ven sensitive (n=7) or ven resistant (n=5). **B.** Principal component analysis was performed on all genes expression data after batch effect was removed using limma and variance stabilized using DESeq2. Resistant samples are in red, and sensitive are in teal. **C.** All genes in the GO biological process Calcium Mediated Signaling pathway were considered for this analysis. Batch corrected and variance stabilized gene expression values for these genes were z-scored and clustered on a heatmap. Black annotation on the left depicts whether or not the gene was considered significant in the DE test (padj < 0.05). Both rows and columns are clustered. **D.** Volcano plot depicting differentially expressed genes. Differential expression analysis was performed on previously published data using DESeq2 comparing sensitive (n = 7) to resistant (n = 5) and controlling for batch effect [41]. Differentially expressed genes were defined as padj < 0.05 and absolute log2FoldChange > 0.5. 1,984 genes were significantly increased in sensitive; whereas 1,953 genes were significantly increased in resistant. **E.** GSEA results in waterfall plot format of all calcium related pathways in the GO biological process pathway set. **F-J.** CITE-seq analysis performed on n=9 ven sensitive primary human AML specimens or n=14 ven resistant primary human AML specimens. **F.** Presto Wilcoxon rank test was used to determine the top 10 markers for each cluster using the log2 fold change values. These markers heavily informed on cell types within each cluster. **G.** Presto Wilcoxon rank test used to determine the top 10 markers for each cluster using the log2 fold change values. These markers heavily informed on cell types within each cluster. Note that some of the genes are duplicated in the matrix because they were called markers for multiple clusters. Expression is z-scored normalized RNA measurement. **H.** Featureplot showing GOBP Calcium Mediated Signaling Module Scores for Normal Bone Marrow Samples (n=3). **I.** Featureplot showing GOBP Calcium Mediated Signaling Module Scores for Sensitive Samples (n=9). **J.** Featureplot showing GOBP Calcium Mediated Signaling Module Scores for Resistant Samples (n=14). **K.** Confocal microscopy to confirm localization of Rhod2AM signal. Representative images from AML #4 and AML#18 that were sorted for ROS-low cells. Green signal is from mitotracker-GFP, red signal is positive signal from Rhod2AM (552/581) while blue signal is from Dapi staining. Individual staining and co-localization staining are shown.

**Supplementary Figure 2.**
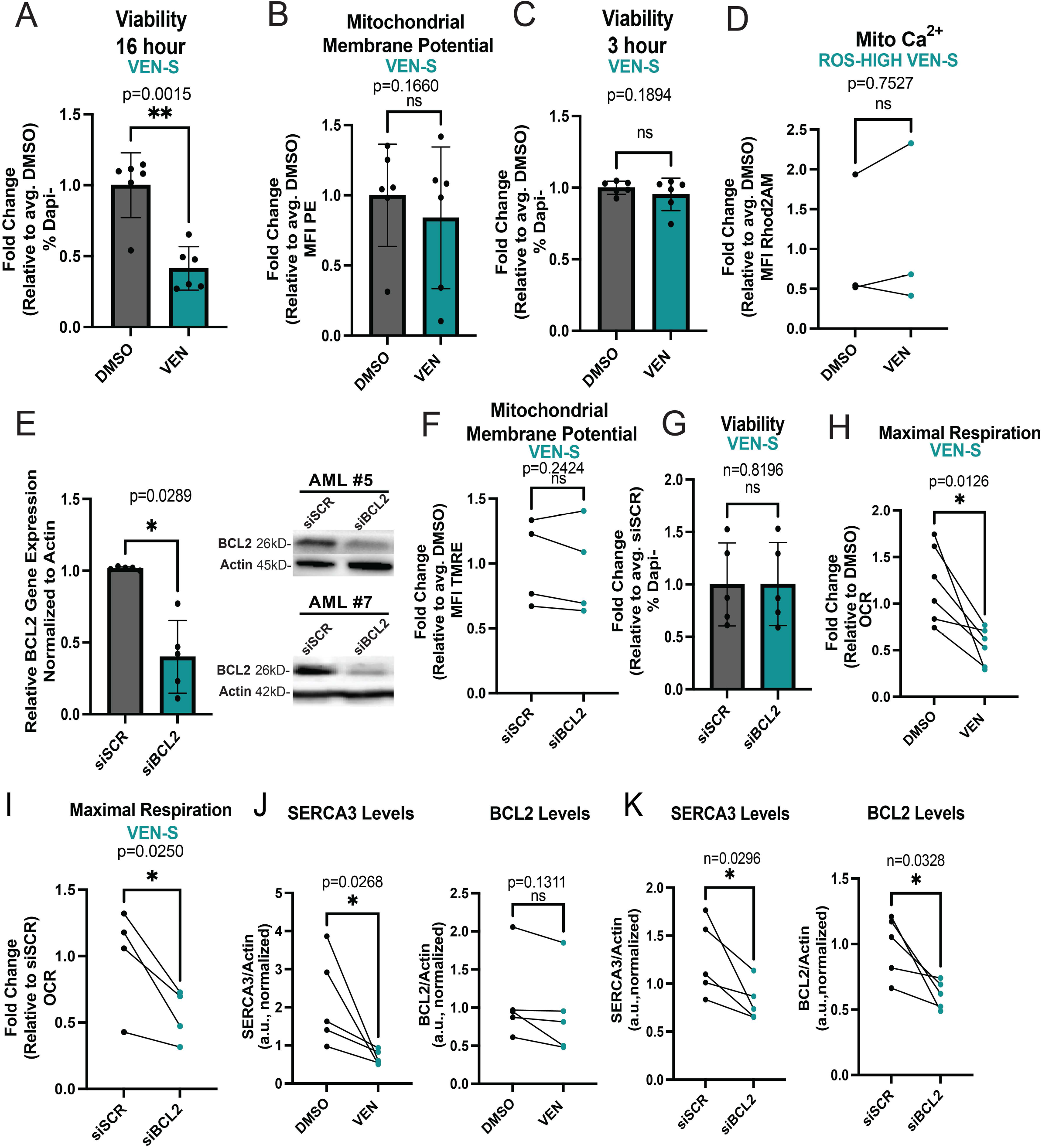
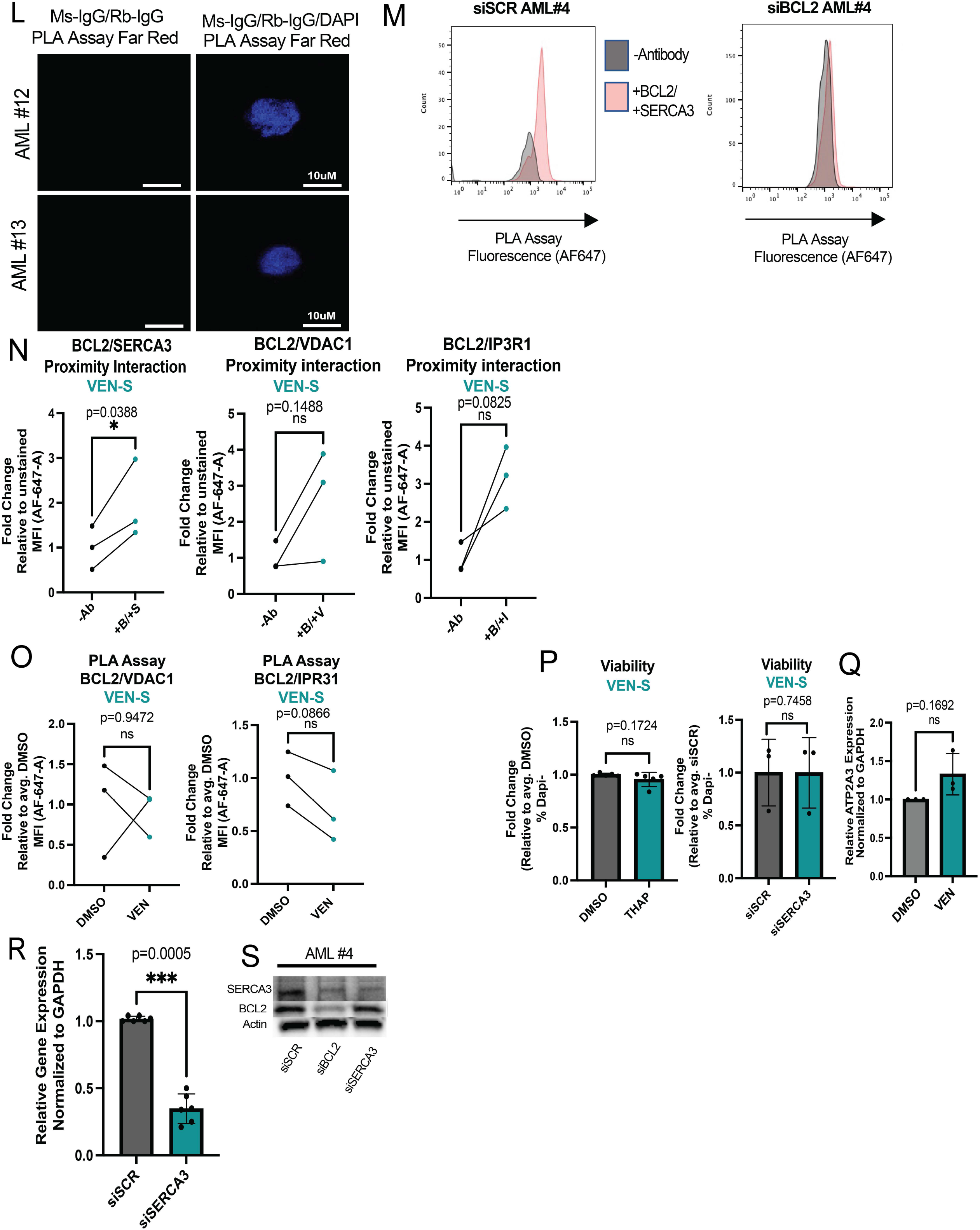
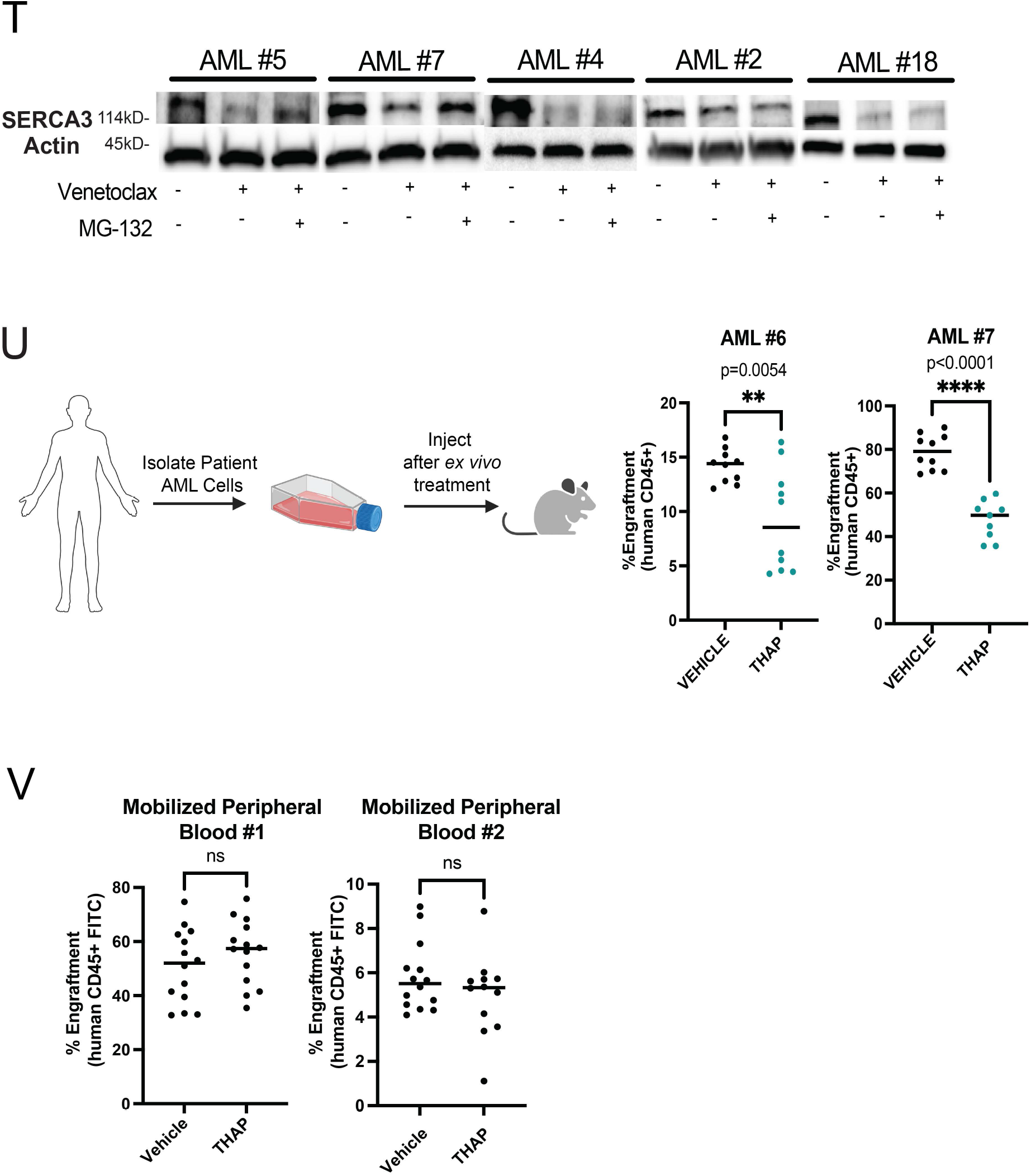
BCL-2 Inhibition Causes Mitochondrial Calcium Changes Associated with SERCA Disruption in Venetoclax Sensitive LSCs. **A.** Viability of venetoclax sensitive primary human AML ROS-low cells after venetoclax treatment (500nM, 16 hours). N=6 (AML# 2,4-7, 18). Data are presented as mean +/-SD. Significance was determined using two-tailed ratio paired t-test. . **B.** Mitochondrial membrane potential of venetoclax sensitive primary human AML ROS-low cells after venetoclax treatment (500nM, 3 hours). N=6 (AML# 2,4-7, 18). Data are presented as mean +/-SD. Significance was determined using two-tailed ratio paired t-test. **C.** Viability of venetoclax sensitive primary human AML ROS-low cells after venetoclax treatment (500nM, 3 hours). N=6 (AML# 2,4-7, 18). Data are presented as mean +/-SD. Significance was determined using two-tailed ratio paired t-test. **D.** Mitochondrial calcium content presented as mean fluorescence intensity (MFI) after venetoclax treatment (500nM, 3 hours) in n=3 venetoclax sensitive primary human AML specimens (AML# 2,4,5) that were sorted for ROS-High cells. Significance was determined using ratio paired t-test **E.** Venetoclax sensitive primary human AML ROS-low cells were electroporated with either siRNA scramble sequence or siRNA for BCL-2 as described in the methods. qRT-PCR was performed to confirm knockdown in AML #2,4-7,18 36 hours post infection. Actin was used as housekeeping gene and expression was normalized to actin expression. Data are presented as mean +/-SD. Significance was determined using two-tailed ratio paired t-test. Western blot to confirm knockdown at 36 hours post infection with representative blots shown for AML #5 and #7. Additional blots showing confirmation of knockdown for AML #2, #4, #18 are shown in Fig. 2E. Anti-actin antibody was used as loading control and anti-BCL-2 antibody was used to determine knockdown. **F.** Mitochondrial membrane potential after BCL-2 knockdown in primary human AML ROS-low cells. N= 4 biological replicates (AML# 2,5,7,18). Data are presented as mean +/-SD. Significance was determined using two-tailed ratio paired t-test. **G.** Viability of primary human AML ROS-low cells (AML #2,4-7,18) after BCL-2 knockdown. Cells were analyzed by flow cytometry 36 hours after infection. Data are presented as mean +/-SD. Significance was determined using two-tailed ratio paired t-test. **H.** OCR after venetoclax treatment (500nM, 3 hours). N=6 venetoclax sensitive primary human AML ROS-low cells (AML# 2,4-7, 18). Significance was determined using two-tailed ratio paired t-test. **I.** OCR after genetic knockdown of BCL-2. N=4 (AML# 2,4,5,7) venetoclax sensitive primary human AML ROS-low cells. Cells were analyzed by flow cytometry 36 hours after infection Significance was determined using two-tailed ratio paired t-test. **J.** Quantification using ImageJ of respective bands from Figure 2D are presented. Significance was determined using two-tailed ratio paired t-test. **K.** Quantification using ImageJ of respective bands from Figure 2E are presented. Significance was determined using two-tailed ratio paired t-test. **L.** Confocal microscopy was used to determine background signal from PLA Assay (Far Red) in primary human AML ROS-low cells. Ms-IgG and Rb-IgG were used for primary antibody incubation steps. Representative images from AML #12 and #13 are presented. **M.** Primary human AML ROS-low cells electroporated with either siRNA scramble sequence or siRNA for BCL-2 as described in the methods. Confirmation of knockdown was assessed at 36 hours post infection as shown in Supplementary 2F. BCL-2 and SERCA3 proximity interaction was measured by PLA assays analyzed through flow cytometry to show specific binding of BCL-2 antibody to BCL-2. **N.** BCL-2 and SERCA3, VDAC1 or IP3R1 proximity interaction as measured by PLA assays presented as MFI. N=3 (AML#2,4,7) venetoclax sensitive primary human AML specimens sorted for ROS-low cells. Significance was determined using two-tailed ratio paired t-test. **O**. BCL-2 and SERCA3, VDAC1 or IP3R1 proximity interaction as measured by PLA assays presented as MFI upon venetoclax treatment (500nM, 3 hours). N=3 (AML#2,4,7) venetoclax sensitive primary human AML specimens sorted for ROS-low cells. Significance was determined using two-tailed ratio paired t-test. **P.** Viability of venetoclax sensitive primary human AML ROS-low cells after thapsigargin treatment (500nM, 3 hours). N=5 (AML# 2,4-7, 18). Data are presented as mean +/-SD. Significance was determined using two-tailed unpaired t-test. Viability of primary human AML ROS-low cells (AML #2,5,7) after SERCA3 knockdown. Cells were analyzed by flow cytometry 36 hours after infection. Data are presented as mean +/-SD. Significance was determined using two-tailed ratio paired t-test. **Q.** qRT-PCR was performed after venetoclax treatment (500 nM, 3 hours) in venetoclax sensitive primary human AML ROS-low cells to determine ATP2A3 expression compared to DMSO treated cells in N=3 (AML# 2,4,7). Actin was used as housekeeping gene and expression was normalized to actin expression. Data are presented as mean +/-SD. Significance was determined using two-tailed ratio paired t-test. **R.** qRT-PCR was performed to confirm knockdown in AML #2, 4-7. Actin was used as housekeeping gene and expression was normalized to actin expression. Data are presented as mean +/-SD. Significance was determined using two-tailed ratio paired t-test. **S.** Representative western blot showing knockdown of SERCA3 in primary human AML specimen sorted for ROS-low cells using siRNA targeting SERCA3 at 36 hours post infection Anti-actin antibody was used as loading control and anti-SERCA3 antibody was used to determine knockdown. **T.** Venetoclax sensitive primary human AML ROS-low cells (n=5) were treated with either DMSO, venetoclax alone (500 nM, 3 hours) or pre-incubated with MG-132 (1uM, 2 hours) then treated with venetoclax (500 nM, 3 hours). Western blot was done to analyze SERCA3 levels in various treatment groups. Anti-actin antibody was used as loading control and anti-SERCA3 antibodies were used to determine protein levels of SERCA3. **U.** Thapsigargin treatment (500nM, 16 hours, ex vivo) and subsequent engraftment potential of venetoclax sensitive primary human AML specimens after transplantation into immune-deficient mice. N=10 for DMSO control group and n=10 and 8, for thapsigargin treatment group for AML 6,7 respectively. 2 million cells injected per mouse and engraftment was assessed between 4-8 weeks. Data are presented as mean with individual data points. Significance was measured by two-tailed unpaired t-test. **V.** Thapsigargin treatment (500nM, 16 hours, ex vivo) and subsequent engraftment potential of normal mobilized peripheral blood samples after transplantation into immune-deficient mice. N=14 for DMSO control group and n=13 and 12, for thapsigargin treatment group for Mob Peri #1 and 2, respectively. 1 million cells injected per mouse and engraftment was assessed between 4-8 weeks. Data are presented as mean with individual data points. Significance was measured by two-tailed unpaired t-test.

**Supplementary Figure 3.**
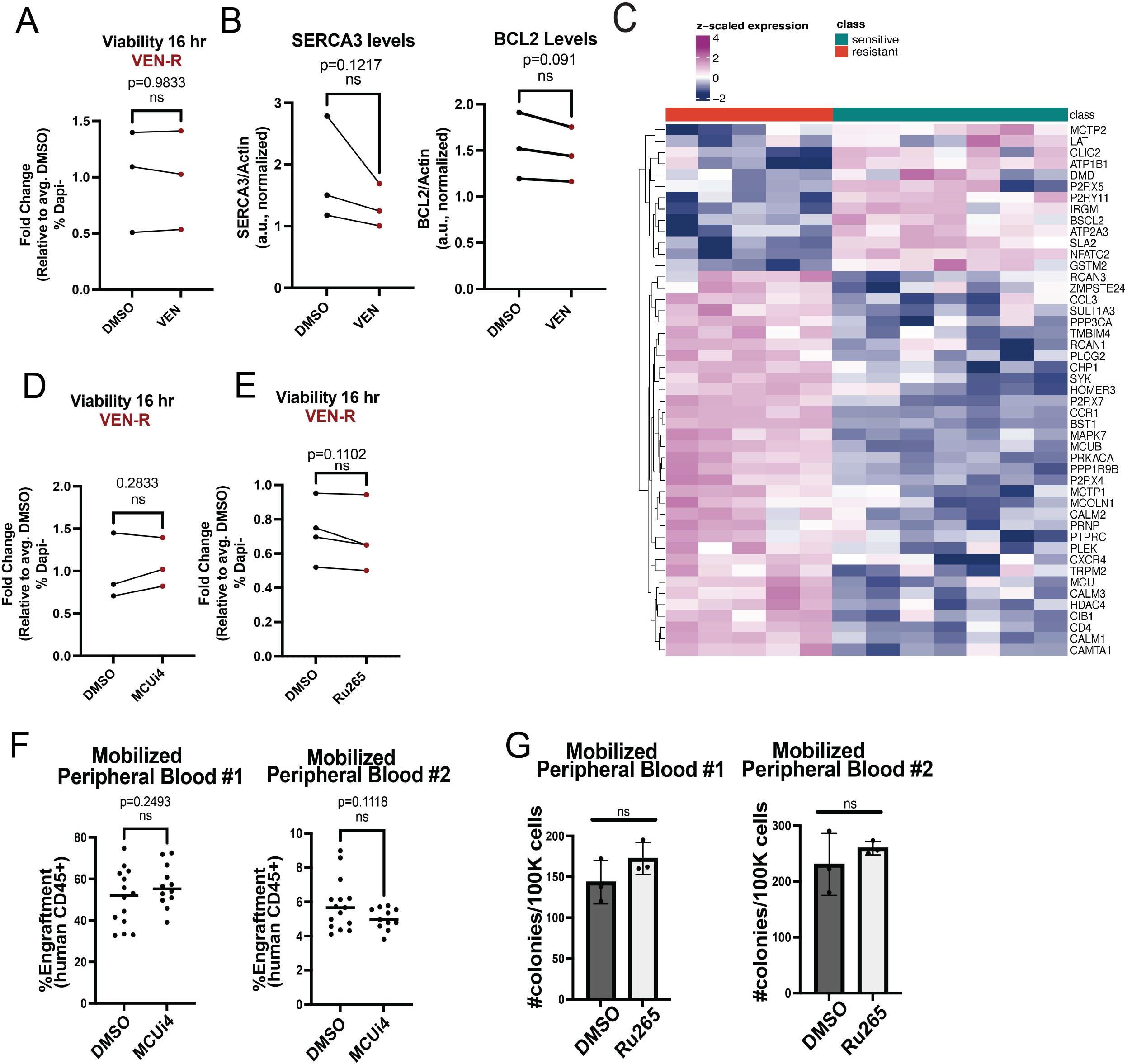
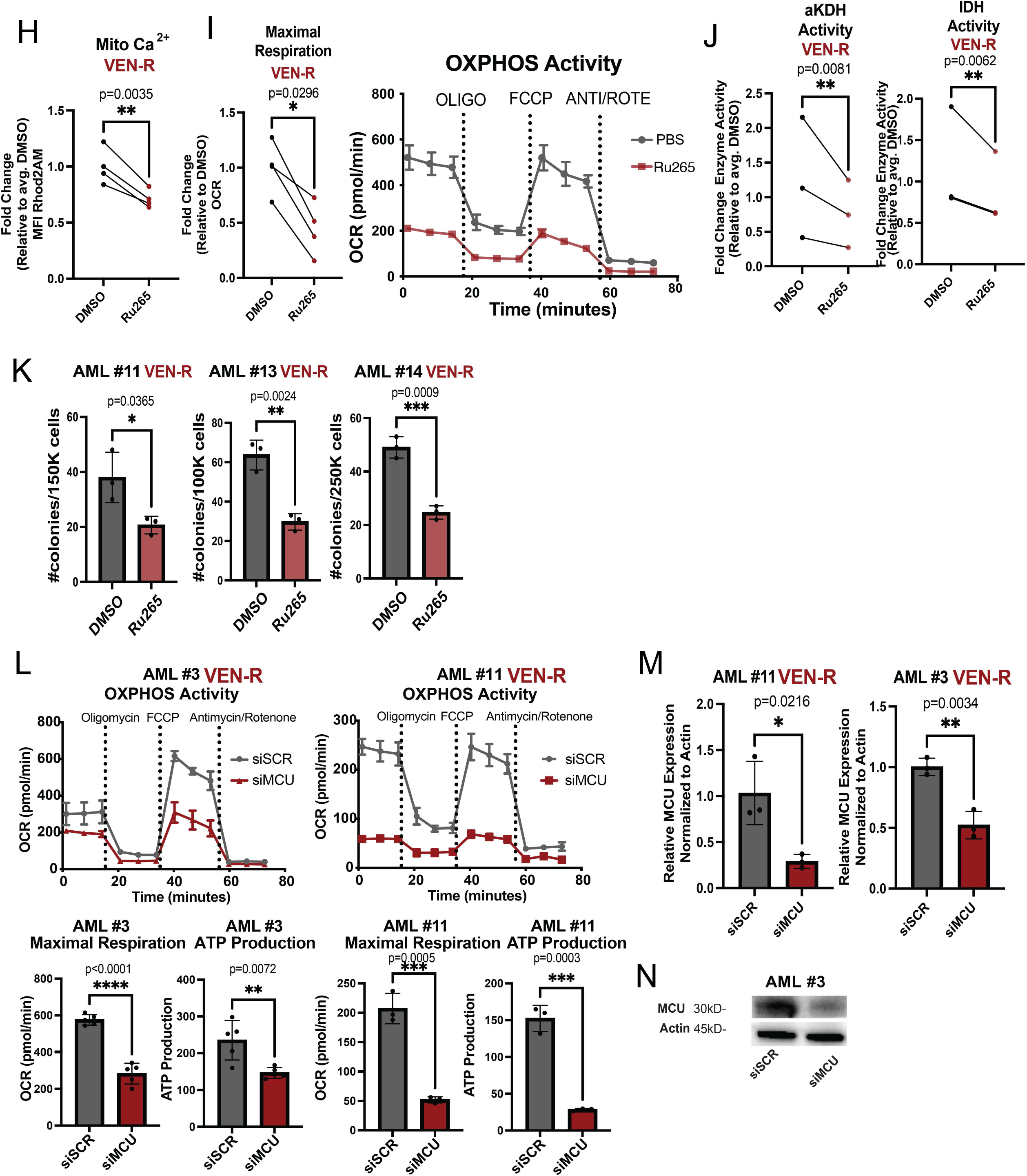
Reducing Mitochondrial Calcium Levels Targets Venetoclax Resistant LSCs. **A.** Viability of venetoclax resistant primary human AML ROS-low cells after venetoclax treatment (500nM, 16 hours). N=3 (AML#11-13). Significance was determined using two-tailed ratio paired t-test. **B.** Quantification using ImageJ of respective bands from Figure 3C are presented. Significance was determined using two-tailed ratio paired t-test. **C.** Only significant (padj < 0.05, absolute log2FoldChange > 0.5) genes in the GO biological process Calcium Mediated Signaling pathway were considered for this analysis. Batch corrected and variance stabilized gene expression values for these genes were z-scored and clustered on a heatmap. Resistant samples are annotated in red, sensitive samples are in teal. Both rows and columns are clustered. **D.** Viability of venetoclax resistant primary human AML ROS-low cells after MCUi4 treatment (5uM, 16 hours). N=3 (AML#11-13). Significance was determined using two-tailed ratio paired t-test. **E.** Viability of venetoclax resistant primary human AML ROS-low cells after Ru265 treatment (10uM, 16 hours). N=3 (AML#11-14). Significance was determined using two-tailed ratio paired t-test. **F.** Engraftment potential of healthy mobilized peripheral blood samples after MCUi4 treatment (5uM, 16 hours, *ex vivo*) after transplantation into immune-deficient mice. Data are presented as mean values with individual data points. Significance was measured by two-tailed unpaired t-test, and n=14 technical replicates for DMSO control group and n=14 and 12 technical replicates for MCUi4 treatment groups for Donor #1 and #2 respectively. **G.** Colony forming units measured by colony formation assays in healthy mobilized peripheral blood samples after Ru265 treatment (10uM, 16 hours). Data are presented as mean values +/-SD. Significance was measured by two-tailed unpaired t-test. N=3 replicates per sample per condition. **H-K.** Venetoclax resistant primary human AML ROS-Low cells (n=4, AML#11-14) treated with Ru265 for 16 hours at 10uM. **H.** Mitochondrial calcium content after Ru265 treatment presented as mean fluorescence intensity (MFI). Significance was determined using two-tailed ratio paired t-test. **I.** OCR after Ru265 treatment. Significance was determined using two-tailed ratio paired t-test. **J.** Isocitrate dehydrogenase activity and alpha keto-glutarate dehydrogenase activity after Ru265 treatment. Significance was determined using two-tailed ratio paired t-test. **K.** Colony forming units measured by colony formation assays in venetoclax resistant primary human AML specimens after Ru265 treatment (DMSO, 10uM Ru265, 16 hour treatment, *ex vivo*). Data are presented as mean values +/-SD. Significance was measured by two-tailed unpaired t-test. N=3 per sample per condition. **L.** OCR after genetic knockdown of MCU in venetoclax resistant primary human AML specimens. N=5 replicates per AML specimen. Significance was determined using two-tailed unpaired t-test. **M.** qRT-PCR was performed to confirm knockdown in AML #3 and #11. Actin was used as housekeeping gene and expression was normalized to actin expression. Data are presented as mean +/-SD. Significance was determined using two-tailed unpaired t-test. **N.** Representative western blot showing knockdown of MCU in venetoclax resistant primary human AML specimen using siRNA targeting MCU at 48 hours post infection Anti-actin antibody was used as loading control and anti-MCU antibody was used to determine knockdown.

**Supplementary Figure 4.**
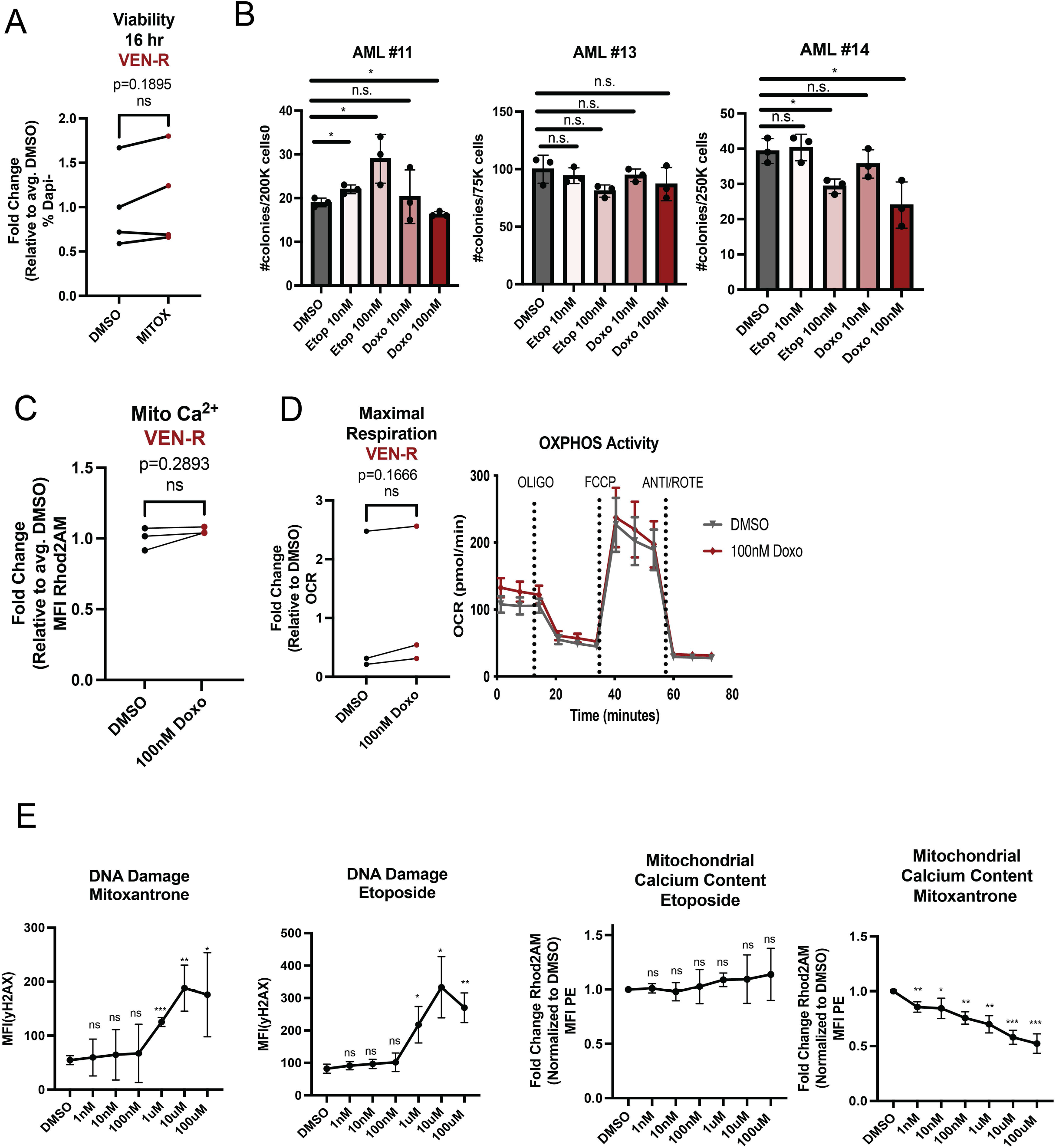
Mitoxantrone Inhibits Mitochondrial Metabolism and Colony Formation in Venetoclax Resistant LSCs. **A.** Viability of venetoclax resistant primary human AML ROS-low cells after venetoclax treatment (100nM, 16 hours). N=4 (AML#11-14). Significance was determined using two-tailed ratio paired t-test. **B.** Colony forming units measured by colony formation assays in venetoclax resistant primary human AML specimens after etoposide or doxorubicin (DMSO,10nM and 100nM, 16 hour treatment, *ex vivo*). Data are presented as mean values +/-SD. Significance was measured by two-tailed unpaired t-test. N=3 per sample per condition. **C and D**. Venetoclax resistant primary human AML ROS-low cells treated with doxorubicin for 16 hours at 100nM. **C.** Mitochondrial calcium content after doxorubicin treatment presented as mean fluorescence intensity (MFI). Significance was determined using two-tailed ratio paired t-test. (N=3, AML#11-13) **D.** OCR after doxorubicin treatment. Significance was determined using two-tailed ratio paired t-test. (N=3, AML#11-13). **E.** Gamma H2AX or mitochondrial calcium content (Rhod2AM) presented as mean fluorescence intensity (MFI) after treatment of MOLM-13 cell line with dose curve of etoposide or mitoxantrone concentrations. Cells were treated with drug for 1 hour due to confounding cell death at higher concentrations at later time points. N=3 replicates.

**Supplementary Figure 5.**
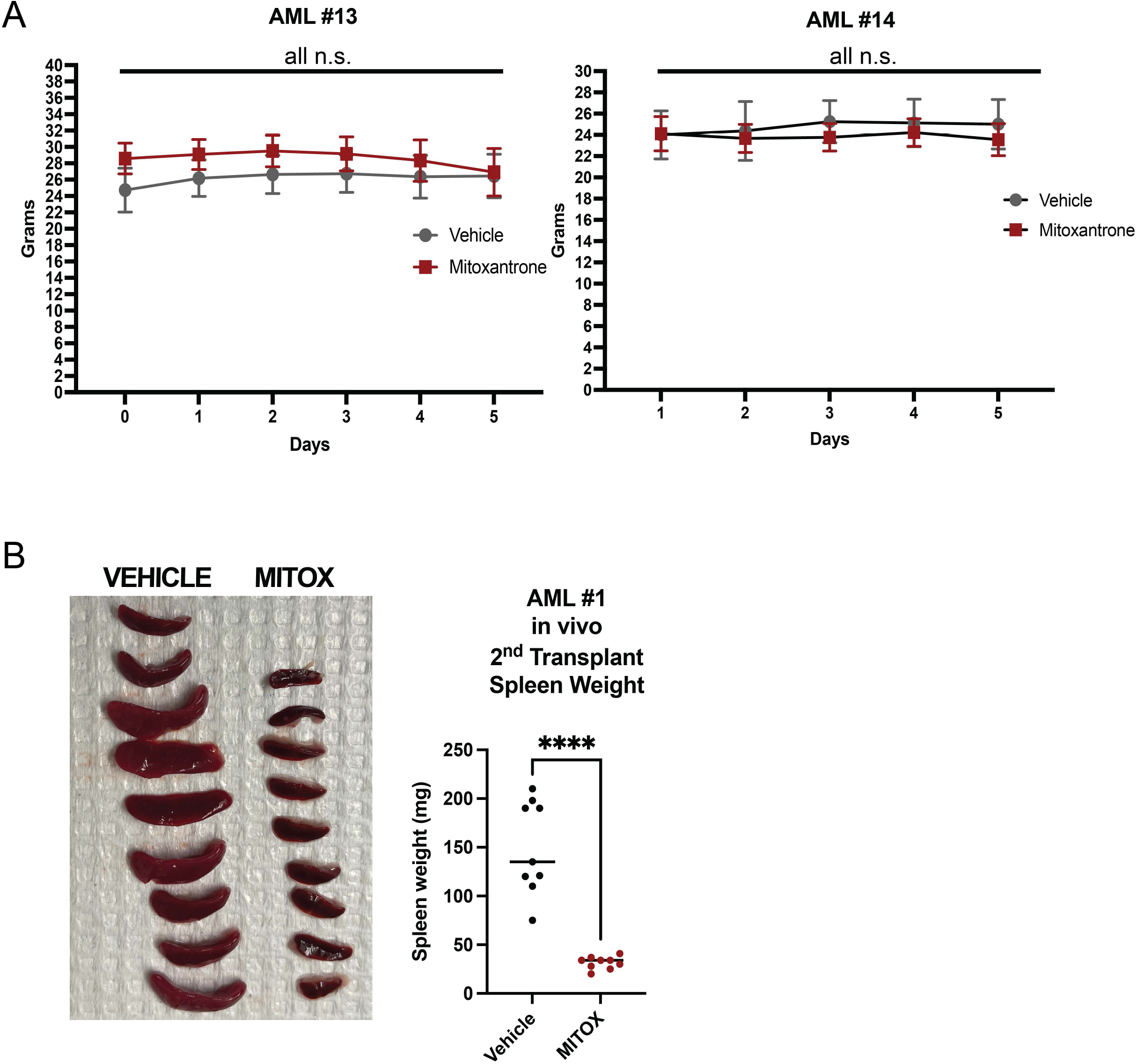
Mitoxantrone Targets Venetoclax Resistant LSCs. **A and B.** Mouse weight average across in vivo vehicle or mitox treatment experiment. N=12 per condition per dose for AML#13 patient sample derived xenograft. N=8 (vehicle), n=9 (mitoxantrone) per condition per dose for AML#14 patient sample derived xenograft. Data are presented as mean values +/-SD. Significance was measured by two-tailed unpaired t-test. **B.** Picture of spleen size of mice harvested after secondary transplantation of cells from Figure 5B (AML #1). Spleen weight in milligrams and data presented as individual values, significance measured by two-tailed unpaired t-test. P<0.0001

